# High-Specificity Detection of Chromosomal Mosaicism Reveals Cell-Type-Specific Genomic Alteration Patterns in Aging Tissues

**DOI:** 10.64898/2026.08.02.741928

**Authors:** Xinyi E. Chen, Hanzhi Wang, Yilin Yang, Marcos G. Teneche, Peter D. Adams, Parker C. Wilson, Nancy R. Zhang

## Abstract

Mosaic chromosomal alterations (mCAs) increase with age and are associated with multiple diseases, yet the cell types and states that harbor these alterations remain largely unknown. Because mCAs arise in individual cells prior to clonal expansion, they are typically rare and obscured in bulk data. We develop CHASM, a method for detecting chromosomal copy number alterations (CNA) from single-cell chromatin accessibility (scATAC-seq) data, a scalable modality that captures both cell state and chromosomal alterations. CHASM estimates a CNA-null background for each cell, providing an individualized expectation for chromosomal accessibility, which is critical in non-neoplastic tissues where alteration-carrying cells are not readily distinguishable from normal. By comparing each cell against its expected background, CHASM distinguishes chromosomal alterations from background variation and achieves more stringent control of false positives. We validate CHASM using in silico spike-in experiments, cross-modality comparisons with matched single-cell DNA and RNA data, and established genome-instability contrasts, including p53 deficiency and chromosome Y loss. Applied to multiple aging data sets, CHASM consistently recovers mCA burden in age-susceptible cell populations and reveals aging-associated signatures not detected by existing methods. In a cohort of 99 human kidney samples spanning age and disease conditions, CHASM identifies enrichment of mCAs in injury-associated cell states (VCAM1-high proximal tubule cells). Notably, CHASM detects the age-associated emergence of mCAs in cancer-relevant genomic regions, including chromosomes 3 gains and losses and chromosome 7 gain, in ostensibly normal cell populations. Cells harboring mCAs exhibit activation of injury-response regulatory programs and reduced epithelial identity programs, while elevated mCA burden in specific epithelial populations are associated with increased immune and stromal infiltration. Overall, we develop CHASM for high-specificity detection of CNA at single cell resolution. Applied across tissues, CHASM reveals aging-patterns of genome instability within cell types and implicates mCAs in early, pre-disease cellular states.

## INTRODUCTION

Somatic mutations accumulate as individuals age due to occasional errors during DNA replication and cell division, which can be influenced by environmental or lifestyle factors^1–4^. Although this phenomenon is well recognized, our understanding of how somatic mutations distribute across cell types with age remain limited. Functionally, somatic mutations may be neutral or deleterious, or they can lead to clonal expansion and malignant transformation^4^. For example, clonal hematopoiesis of indeterminate potential (CHIP), a common age-related somatic mutation phenomenon arising from hematopoietic stem cells, has been associated with increased risk for blood malignancies, cardiovascular diseases, and other age-associated conditions^5–9^. While somatic mutations in blood have been better characterized owing to the relative ease of sampling and the frequent occurrence of clonal expansions, little is known about how somatic mutations are distributed across cells both in blood and solid tissues.

One class of somatic mutations is chromosomal copy number alterations (CNA), which are characterized by large gains and losses of a chromosome. When such alterations are present in only a subset of cells within a tissue, they are referred to as mosaic chromosomal copy number alterations (mCAs)^1,3,10–12^. Compared with single-nucleotide variants, mCAs are rarer in healthy tissues, yet they tend to associate with disorders^1,3,10^, likely because these large-scale alterations have more pervasive impact on cellular function and may reflect dysfunction in DNA replication and repair machinery. Biobank-scale analyses of blood show that certain mCAs can be markedly enriched^6–8,11–14^. Sex chromosome mCAs, such as mosaic loss of the X or Y chromosome (mLOX and mLOY), are prevalent in blood, with mLOY occurring in more than 30% of males over the age of 70 and mLOX occurring in about 12% of females^15,16^. A pan-tissue study based on bulk RNA-seq data demonstrate that mCA burden varies across tissues, with esophagus and adrenal glands exhibiting the highest incidence rate (∼10% individuals) whereas heart and kidney are among the lowest^3^. Despite these broad insights, estimates of mCA frequency across cellular populations have largely been inferred from bulk sequencing data or SNP arrays, which have limited power to detect rare, non-clonal events^1–3,7,11–15^. These approaches offer limited resolution for mCAs on the single cell or cell type level, hindering efforts to characterize the specific cell types in which mCAs arise and consequently to understand their disease relevance.

Understanding how mCAs and genome instability manifest at the level of specific cell types is critical for elucidating its role in aging and disease. Different cell types may vary substantially in their susceptibility to genomic alterations and in their capacity to tolerate or respond to such changes, with potential consequences for tissue dysfunction and chronic disease progression. Bulk tissue measures average signals across heterogeneous cell subsets and do not resolve the cellular contexts in which mCAs arise. As a result, they cannot capture the relationship between genome instability and cell state, nor identify the specific cell populations that may drive age-related pathology. Addressing these questions requires methods that can quantify genome instability and link it to molecular phenotypes at single-cell resolution.

The advent of single-cell sequencing presents an opportunity to characterize mCAs in a cell-type specific manner. Recent co-assay technologies that jointly profile single cell DNA and molecular modalities, such as chromatin accessibility (ATAC) or gene expression^17,18^, enable simultaneous capture of copy number variation and cell type identity, but these approaches remain costly and difficult to scale. As a result, there remains growing interest in inferring copy number alterations directly from scATAC-seq or scRNA-seq data. Existing tools for this purpose have largely been developed in the context of cancer^19–21^, where altered cells are clonally expanded and exhibit distinct features that allow them to be separated from non-malignant cells. In such settings, dominant CNA signals can often be recovered even in the presence of substantial background variation. By contrast, in aging tissues, CNAs occur in much lower frequencies and there is no clear way to identify normal cells, both of which challenge the existing frameworks for CNA detection. In particular, because true CNAs in these tissues are rare and may not be clonally expanded, their detection requires stringent control of false positives. These challenges limit the applicability of existing approaches and highlight the need for methods that can robustly distinguish low-frequency CNA signals from pervasive background variation in single-cell sequencing data.

Single-cell ATAC-seq, in particular, provides a powerful modality for studying CNA in aging tissues, as it captures chromatin accessibility patterns that reflect cell state while allowing for the computation of epigenetic aging-related features^22,23^. Widely adopted co-assay technologies now enable joint profiling of chromatin accessibility and gene expression at single-cell resolution, and large-scale scATAC-seq datasets have been generated across diverse aging tissues^24,25^. In this study, we develop CHromosomal Alterations in Somatic Mosaicism (CHASM), a framework for robust detection of CNAs using single-cell ATAC data, with particular emphasis on achieving the level of precision required for confident detection of low-frequency events. We rigorously benchmark CHASM using controlled spike-in experiments in paired scDNA–scATAC data to assess sensitivity and specificity, and further validate via matched scDNA and scRNA data, as well as contrasts between cell populations with established differences in genome instability. These benchmarks and validations demonstrate that CHASM achieves the robustness and precision required for mCA profiling in tissue aging studies.

We then apply CHASM across multiple aging datasets to ask how chromosomal mosaicism accumulates across tissues and cell types. In young and aged hematopoietic stem cells, CHASM detects increased CNA burden with age, consistent with the established accumulation of chromosomal mosaicism in the hematopoietic system^11–14^. Across a whole-organism mouse aging atlas^26^, CHASM-detected CNA burden shows age-associated increases in a subset of tissues and annotated cell types, with signals more frequently observed in epithelial populations. In aging liver multiome data, increased CNA burden in hepatocytes is further associated with transcriptional disorder, linking ATAC-inferred chromosomal instability to an independent RNA-derived measure of cellular dysregulation. Together, these analyses show that CHASM can recover coherent aging-associated patterns across tissues while revealing cell type–specific susceptibility to chromosomal instability.

We further apply CHASM to characterize the cell type specific distribution of mCAs in kidney tissue across 99 human samples spanning age and disease conditions. We find that VCAM1-high injured proximal tubule cells harbor markedly elevated CNA burden. Beyond this injured population, CNA burden increases with age among cells that appear otherwise unremarkable, with preferential enrichment in proximal tubule and thick ascending limb cells. Cells with higher CNA burden exhibit elevated expression of injury-related genes and transcription factor binding activity. Increased CNA burden is further associated with greater infiltration of immune and stromal cell within the tissue microenvironment. Age-associated increases in mCA preferentially localize to specific genome regions, several of which are recurrently altered in kidney cancer. Together, these findings indicate that genome instability emerges in specific epithelial compartments during human aging and is linked to early signals of injury and tissue remodeling, even in the absence of overt disease.

## RESULTS

### Overview of the CHASM algorithm

Our goal is to detect copy number alterations (CNAs) in scATAC-seq data from normal, non-dysplastic tissues, where cells harboring these alterations have not undergone clonal expansion and are therefore low frequency. Identifying such rare signals requires high specificity to reliably distinguish true aberrant cells from background variation. Otherwise, true positive CNAs can be easily obscured by false positives, making it more difficult to draw accurate biological conclusions. The central challenge in scATAC-based CNA detection is that large-scale variation in chromatin accessibility, driven by cell state and technical effects, can mimic the broad shifts expected from copy number alterations, leading to false positives if not properly removed. To illustrate this, we first examine a paired scDNA and scATAC dataset from healthy breast tissue^17^, where we use the DNA modality to filter for cells that do not carry any CNA. For these confident “true negative” cells, we bin read counts along each chromosome, normalize by total library size, and hierarchically cluster based on their normalized bin count profile (**Fig. 1A, Methods**). This results in three main clusters, each exhibiting strikingly broad mean shifts along the genome that could be mistaken as copy number variation. The three clusters are loosely associated with the three cell types in the dataset, suggesting that some, but not all, of this variation is driven by cell type. Importantly, these differences also persist at the chromosome level, thus smoothing across broad ranges cannot remove cell-state differences. This observation motivates a method that can account for background variation in a dataset adaptative manner.

**Figure 1.**
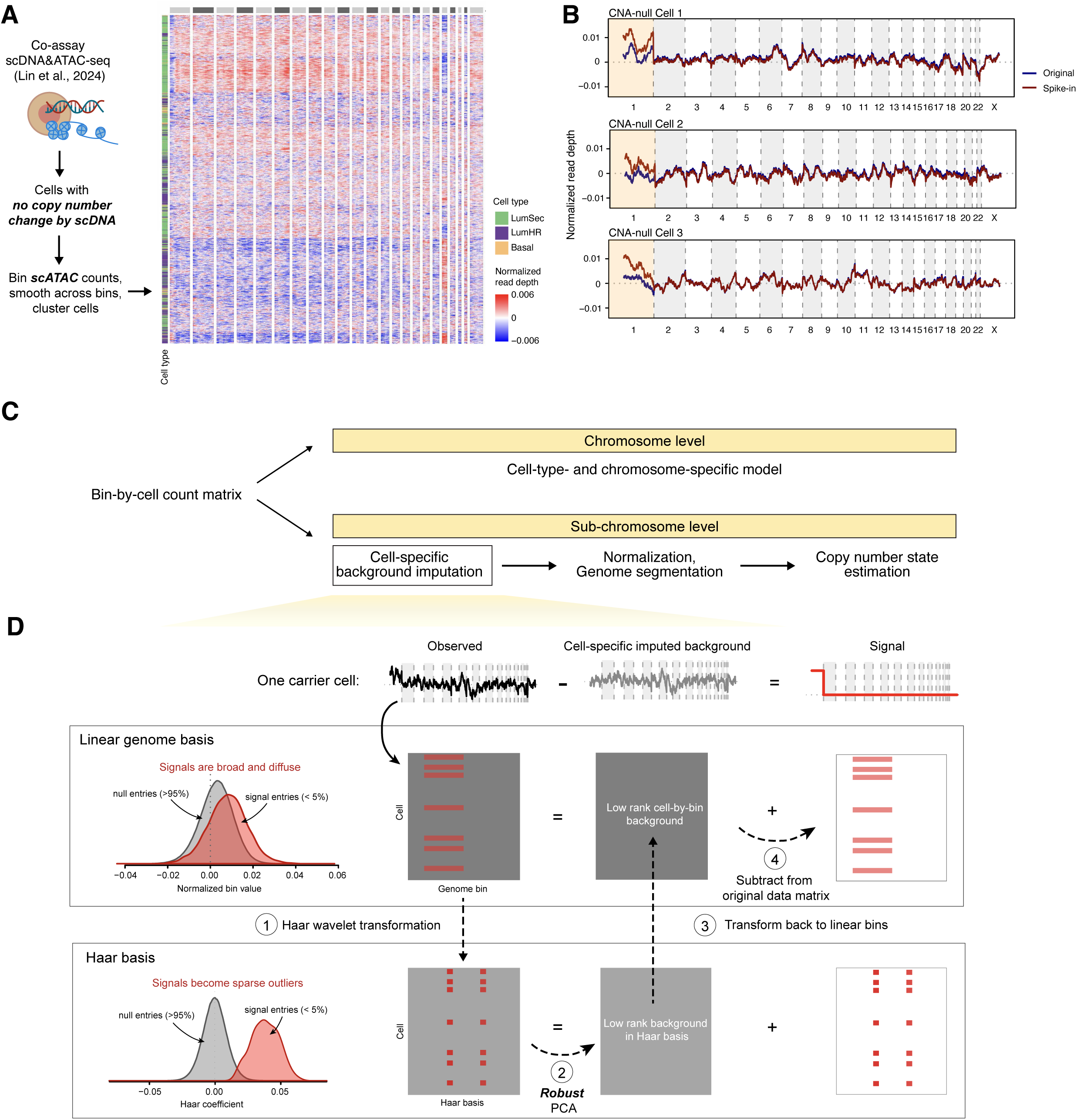
Challenges in modeling copy number states in scATAC-seq data and overview of CHASM algorithm. **A)** Epithelial cells from breast tissues^17^ with no copy number alteration detected from scDNA-seq are used to examine their scATAC read depth profiles. Heatmap shows normalized and smoothed within chromosome ATAC read count profiles clustered by profile similarity. The annotation bar on the left is colored by cell type identity. The annotation bar on the top is separated by chromosome. **B)** Normalized and chromosome-smoothed scATAC-seq read count per genomic bin in three representative cells. Navy blue line shows the original normalized read depth. Dark red line shows the read depth profile with one-copy gain spike-in to chromosome 1. Light orange background marks the spike-in region. **C)** Workflow of the CHASM algorithm. For chromosome-level detection, a cell-type- and chromosome-specific negative binomial distribution approach is used to identify outliers in observed read depth. For sub-chromosomal-level detection, cell-specific background is first estimated, followed by genome segmentation and copy number state estimation. **D)** Illustration of cell-specific background estimation. Library-size normalized read count profiles are projected from the linear genome to Haar wavelet basis, followed by robust PCA to separate background variations and outlier signals. Background variation is then transformed back to the linear genome scale, and this represents the cell-matched control normalized read count matrix. The control profile is then subtracted from the observed data to obtain a signal residual matrix. Genome segmentation is performed on the residual matrix to obtain break points. Copy number states are assigned by comparing the observed and imputed background read depth profiles.

What would real copy number changes look like amid this background? We spiked in a one copy gain in chromosome 1 in a subset of cells and performed read depth normalization as before (**Methods**). Despite the spike-in being applied to the whole of the largest chromosome, its distinction is not obvious against the background fluctuations (**Fig. 1B**). As shown in Figure 1A and 1B, these background variations are cell- and loci-specific, motivating a method to estimate a **cell-specific “no-CNA” baseline** directly from the data.

First, we focus on a strategy for the simpler case of whole chromosomal aneuploidies. Aggregating reads across a full chromosome largely mitigates the effects of data sparsity and cell state differences in scATAC data. We use a negative binomial distribution to model chromosomal read count in a cell type and chromosome-specific manner (**Fig. 1C**, **Methods**). For each chromosome *r*, the expected read count in a CNA-negative cell is estimated from the median fraction of total reads mapping to *r* among cells of the same type. We then compare the observed read count against the expected by a negative binomial test to determine whether the chromosome-level read depth deviates significantly from the population median. The dispersion in the negative binomial distribution is estimated via a nonparametric strategy. This approach produces a p-value for each chromosome in each cell, enabling explicitly control of false discovery.

Next, consider the more difficult case of sub-chromosomal alterations, for which we construct a cell-specific reference. To address data sparsity, we aggregate reads into fixed-width genomic bins (width depending on data quality, see Methods). We reason that CNAs create relatively rare and distinct breakpoints in read depth along a chromosome. These breakpoints are more easily detectable after Haar wavelet transformation, a signal processing technique that separate local, abrupt shifts from background fluctuations (**Fig. 1C, D**, **Methods**). This step converts broad shifts in mean along genomic positions, corresponding to copy number signals, into sparse, high-magnitude coefficients in the Haar wavelet basis.

We further assume that background variation can be captured by a relatively low number of background latent factors due to cell type or other technical batch effects. Since the Haar transformation is linear, it does not disrupt this low-rank structure. Thus, the cell-by-bin wavelet transformed matrix can be decomposed into a sum of two parts: a low-rank background and a copy number signal that is sparse and high magnitude. This enables us to use robust PCA to separate the low-rank component, which captures the systemic background variation, from the residual component, which captures the sparse signal. In other words, because wavelet transformation reduces CNA signals to sparse outliers, the low-rank component from robust PCA is expected to mainly reflect cell type and any other shared background variations. Thus, after transformation back into the linear genome basis, the low-rank component provides a **per-cell expected profile in the absence of CNA**, serving as a normalization reference. Subtracting this reference from the observed data yields a residual component enriched for CNA signals. We perform linear segmentation on this residual matrix to identify candidate CNA regions and assign copy number states by comparing the observed read depth with the estimated cell-specific reference. This step tackles cell-type effects in scATAC data such that normal variations are less likely to be mistaken for CNA signals.

Together, our framework improves scATAC-based CNA detection specificity by modeling chromosome-level read counts with a negative binomial distribution to account for expected variations, and by performing wavelet transformation and robust PCA on the sub-chromosomal level. Wavelet transformation enhances the separation of CNA signals from background and robust PCA accounts for cell-type and chromosome specific variations.

### Validation of CNA detection using scATAC data with spike-in ground truth

We first assess the performance of CNA detection by evaluating both specificity and sensitivity via in silico spike-ins. Specificity refers to the absence of CNA calls in CNA-negative cells and regions, whereas sensitivity refers to accurate detection of CNA in true CNA-positive cells. Our goal is to achieve high sensitivity while controlling false positive detections. Although both metrics matter, specificity has been underexplored in existing scATAC-based CNA detection methods. Given the intended application of our method to nondysplastic tissues where high specificity is essential, we therefore place particular emphasis on demonstrating robust specificity while preserving strong sensitivity.

Our first set of benchmarks make use of the normal breast tissue dataset with paired scDNA and scATAC shown in Figure 1A, B^17^, pre-screened for “true negative” cells that do not carry any CNA as indicated by the matched scDNA modality. As before, we spike-in CNA to the scATAC modality by adding or removing reads from specific genomic regions (**Fig. 2A, Methods**). To comprehensively test the algorithm across event sizes, frequencies, and amplitudes, we design spike-ins to the entire chromosome 1, chromosome 1p, and a 50mb region in chromosome 1. These events are introduced into 1, 5, and 10% of cells, from two copies loss to two copies gain.

**Figure 2.**
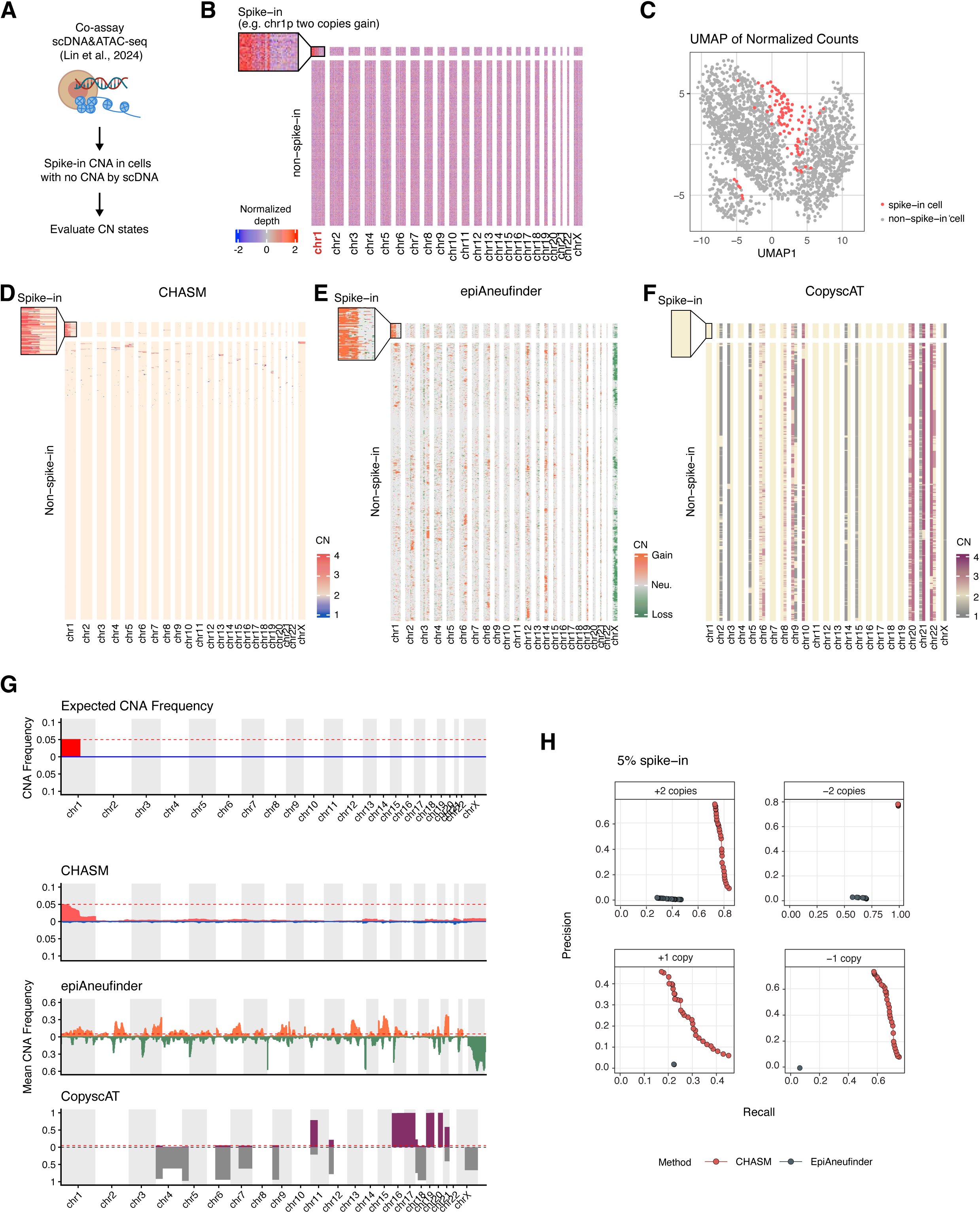
Benchmarking of CNA detection performance using semi-synthetic spike-in data. **A)** Schematic of spike-in experiment on scATAC-seq data on cells with no copy number alteration by scDNA-seq. **B)** Heatmap of normalized read depth. Top panel shows 5% of cells with chromosome 1p two copies gain spike-in. Bottom panel shows the non-spike-in cells. **C)** UMAP visualization generated from normalized read counts. Color labels spike-in status. **D)** Heatmap of copy number state determined by CHASM. **E)** Heatmap of gain or loss status determined by epiAneufinder. **F)** Heatmap of copy number state determined by CopyscAT. **G)** Bar plot of copy number alteration frequency along the genome. Positive direction summarizes copy number gain. Negative direction summarizes copy number loss. The top panel shows the expected CNA frequency based on the design of the spike-in experiment. The bottom three panels are results from CHASM, epiAneufinder, and CopyscAT, respectively. Three replicates are used in this analysis, and the result plotted is the average frequency across replicates. The red dotted line indicates the expected frequency of the copy number alteration event. The color of the bar plots corresponds with gain or loss status. **H)** Pareto frontier of bin-level precision and recall for CNA detection by sweeping the segmentation stringency parameter in CHASM and epiAneufinder. Results are shown for spike-in conditions ranging from two-copy gains to two-copy losses on chromosome 1p and a 50 Mb region of chromosome 1 in 5% of cells.

We compare CHASM to scATAC aneuploidy detection methods epiAneufinder^19^ and CopyscAT^20^. Briefly, epiAneufinder aggregates reads into genomic bins, corrects for GC content, and segments the GC-corrected profile^19^. It then defines a genome-wide baseline using segments with near-average read depth and assigns copy number states against this baseline^19^. CopyscAT aggregates reads into chromosomal arms, generates pseudodiploid controls based on the population median, and applies Gaussian decomposition to identify clusters of CNAs in each chromosome arm^20^.

To illustrate the result, we show examples of two-copy-gain spike-in on chromosome 1p at 5% cells (**Fig. 2B-G)**, and the result of the other conditions can be found in the supplemental figures (**Supplemental Fig. 1-3**). Our method recapitulates the expected spike-in CNAs without introducing large amounts of false detections (**Fig. 2D**). A summary of CNA frequency along the genome by CHASM reveals a clear pattern of the true CNA spike-in, similar to the expected CNA frequency plot (**Fig. 2G**). In comparison, epiAneufinder can recover the spike-in CNAs, but it makes spurious detections at high frequency (**Fig. 2E**). As a result, a summary of CNA events along the genome misleadingly suggests wide-spread CNA patterns, overwhelming the genuine spike-in chromosome 1p gain (**Fig. 2G**). This is likely due to epiAneufinder’s reliance on a genome-wide baseline, which may not adequately account for inherent differences in accessibility along the genome. Consequently, regions with systematically lower accessibility can be misinterpreted as copy number loss. Finally, CopyscAT cannot detect the spike-ins signals while simultaneously reporting widespread alterations in other genome regions (**Fig. 2F, G**), likely due to insufficient normalization for cell-type-specific accessibility differences, as illustrated in Figure 1A.

We next evaluate the precision and recall of spike-in CNA. Different methods provide distinct tuning parameters (“knobs”) that control the trade-off between sensitivity and specificity; we vary these method-specific parameters to trace out the precision–recall relationship. For CHASM, we vary the threshold controlling segmentation stringency (**Methods**). For epiAneufinder, we vary the threshold for which a segment is retained. CopyscAT is excluded here as segmentation is not part of its pipeline, making it not directly comparable in this parameter sweep. As such, this benchmark reflects a sweep over method-specific hyperparameter rather than a standard operation curve. Our method consistently performs better under various thresholds used in segmentation (**Fig. 2H, Supplemental Fig. 4A-C**). Across all conditions, CHASM controls false detection of CNA better than existing tools and achieves much improved specificity without losing the sensitivity for recapitulating the spike-in ground truth.

Finally, to illustrate how the combination of wavelet transformation and robust PCA improves signal separation, we compare t-statistics that quantify the signal-to-noise ratio across CNA breakpoints. Note that in this comparison, we vary only the normalization step of the pipeline, fixing the segmentation and copy number state assignment steps that follow. As a baseline for comparison, we considered simple read depth, or read depth normalized by background estimated via robust PCA alone (see **Methods**). Compared to read depth and robust PCA alone, wavelet transformation combined with robust PCA significantly increased the differences between CNA segment and the surrounding normal regions (**Supplemental Fig. 4D,E**). Robust PCA alone underperforms because true CNAs represent consecutive changes in genomic space rather than sparse signals, making them more likely to be captured by the low-rank component of robust PCA, thus mistaken as background and erroneously removed during normalization. This comparison establishes the unique effectiveness of pairing Haar wavelet transformation with robust PCA in separating signal from background noise.

### Validation of CNA detection using cross-modality data and biological contrasts

We next apply CHASM on real world scATAC-seq data across diverse settings and validate detected CNAs using orthogonal evidence, either through matched cross-modal measurements or established contrasts between cell subsets with known differences in genome instability. For cross-modal validation, we utilize the paired scDNA and scATAC co-assay data on normal breast tissue samples^17^, treating scDNA as the reference standard, as well as scMultiome with paired scRNA and scATAC data^27^, using cell-matched RNA expression as an orthogonal surrogate. As for established biological contrasts, we reason that there should be CNA burden differences between cells with and without Y chromosome loss, a known marker of genome instability^27–30^, and cells with intact or deficient p53 function, given p53’s central role in maintaining genome integrity^31,32^.

First, using the normal breast tissue data from Lin et al.^17^, we compare the mCA profile detected from scDNA and scATAC (**Fig. 3A**). In this dataset, the epithelial component is composed of three cell types: luminal secretory / progenitor cells (LumSec), luminal hormone receptor cells (LumHR), and basal-myoepithelial cells (Basal) (**Supplemental Fig. 5A**). Concordant with Lin et al, the LumSec population harbor a minor population of cells with chr1q gain and chr 10 loss in the scDNA data (**Fig. 3B**). Consistently, CHASM detects about 2.5% of LumSec cells harboring chromosome 1q gain, and about 1.5% of the population with chromosome 10 loss. In contrast, even though both epiAneufinder and CopyscAT detect instances of chromosome 1q gain and chromosome 10 loss, there are pervasive spurious CNA calls not validated by the cell-matched scDNA, obscuring the genuine CNAs.

**Figure 3.**
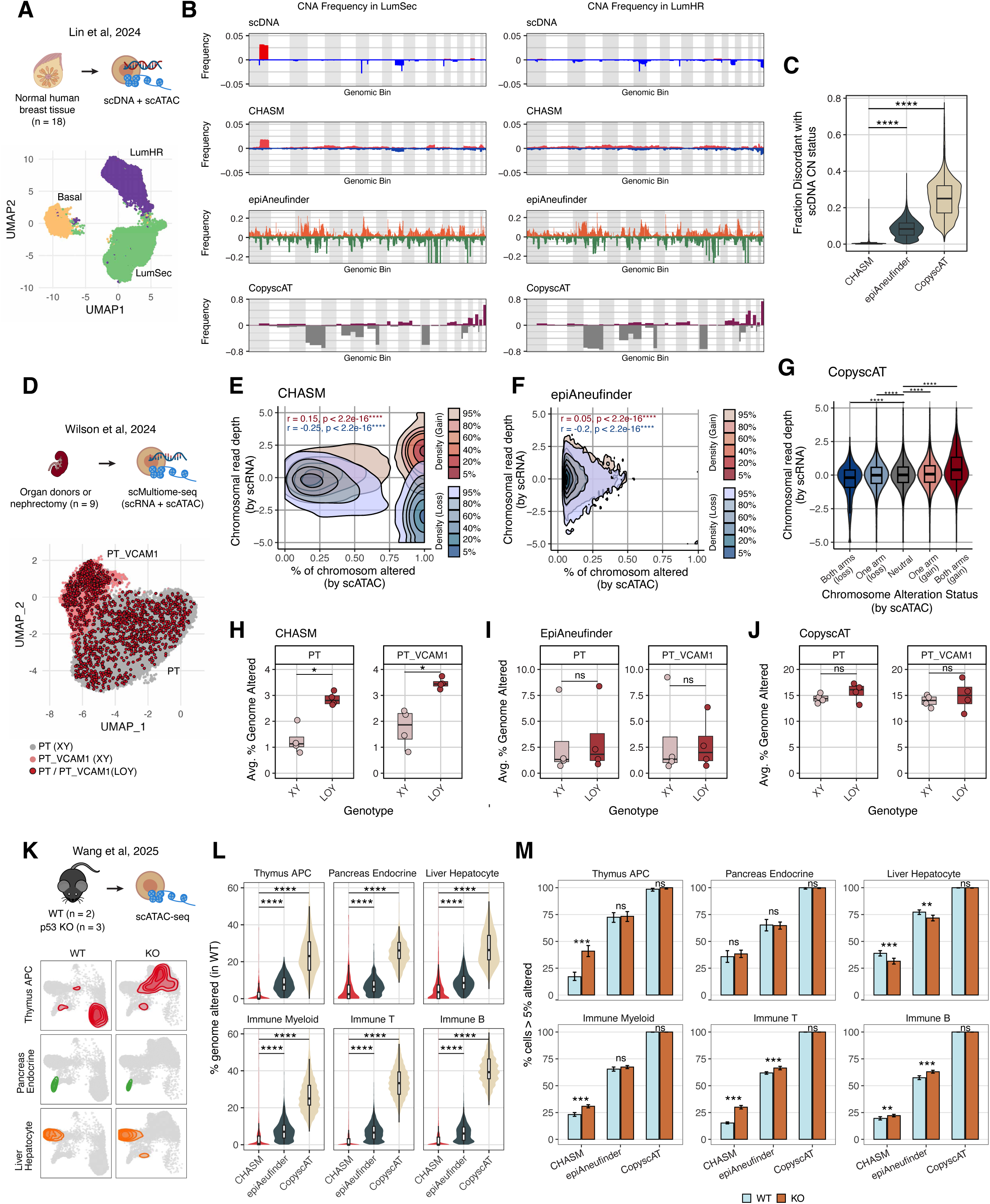
Benchmarking of CNA detection performance using cross-modality data and biological contrasts. **A)** Schematic of normal breast tissue data and UMAP visualization of the scATAC-seq data^17^. **B)** Bar plot of copy number alteration frequency in luminal secretory progenitors (LumSec) and luminal hormone receptor cells (LumHR) along the genome, comparing detection on scDNA-seq and scATAC-seq determined by CHASM, epiAneufinder, and CopyscAT. **C)** Violin plots of proportion of genome with discordant copy number state estimate in scDNA-seq and scATAC-inferred by CHASM, epiAneufinder, and CopyscAT. **D)** Schematic of scMultiome-seq experiment from human kidney tissue^27^. UMAP of proximal tubule (PT) cells and injured proximal tubule cells (PT_VCAM1). Gary and pink indicate PT or PT_VCAM1 cells with normal sex chromosome genotype (XX or XY); red indicates PT or PT_VCAM1 cells with loss of Y chromosome (LOY). **E)** Density contour plot comparing the fraction of chromosome affected by copy number alteration (CNA) inferred from scATAC-seq by CHASM and normalized chromosomal transcript abundance from scRNA-seq. Red colors indicate copy gains, and blue colors indicate copy losses. Spearman correlations are annotated. The highest density regions at 95%, 80%, 60%, 40%, 20%, and 5% are plotted. **F)** Density contour plot comparing the fraction of chromosome affected by CNA inferred from scATAC-seq by epiAneufinder and normalized chromosomal transcript abundance from scRNA-seq. **G)** Violin plots of normalized chromosomal transcript abundance from scRNA-seq in cells with full chromosomal loss, one-arm loss, copy neutral, one-arm gain, and full chromosomal gain determined by CopyscAT. Two-sided Wilcoxon tests are performed against chromosomes with copy number of 2. **H-J)** Box plots comparing the average percentage of genome with CNA determined by CHASM, epiAneufinder, and CopyscAT between cells of normal sex chromosome genotype and LOY genotype in PT and PT_VCAM1 cells. Each dot is an individual. Two-sided Wilcoxon tests are performed. **K)** Schematic of p53 knock out mouse data and UMAP visualization of the scATAC-seq data, highlighting thymus antigen presenting cells (APC), pancreas endocrine cells, and liver hepatocytes. **L)** Violin plots of per-cell percentage of genome with CNA on wild type cells by different tools in thymus APC, pancreas endocrine cells, liver hepatocytes, and immune cells. Two-sided Wilcoxon tests are performed. **M)** Bar plots on the fraction of cells with >5% of genome altered in wild type or p53 knock out conditions based on CHASM, epiAneufinder, and CopyscAT results. Color indicates genotype status. P-values are derived from Chi-squared test.

Additionally, a small subset of the LumHR population harbor chromosome 7, 16, and 22, loss based on scDNA. CHASM captures a subset of chromone 22 loss more evidently but not the other CNA. In comparison, epiAneufinder and CopyscAT report pervasive CNA calls in other regions which are difficult to distinguish from genuine signals. Thus, although CHASM identifies a subset of the CNAs detected by scDNA, its reported events show higher concordance with the matched DNA modality and fewer false positives than existing methods. At the single-cell level, CHASM also show significantly lower discordance in copy number state across the genome between scATAC and scDNA than the other two methods (**Fig. 3C, Supplemental Fig. 5B**).

Next, we turn to a single cell multiome dataset from kidney epithelial tissue (**Fig. 3D**). Wilson et al^27^ reports that a subset of kidney proximal tubule cells (PT) harbor loss of Y chromosome (LOY), particularly in the injured PT cells (PT_VCAM1). To benchmark copy number estimation reliability, we first compare to read count from cell-matched RNA. We reason that true CNAs are likely accompanied by concordant shifts in RNA abundance, with copy number gains associated with increased expression and copy number losses associated with decreased expression. Comparing the fraction of chromosome altered reported in scATAC to scaled chromosome transcript reads in scRNA, we observe a significant correlation (r = 0.15 and −0.25 for CN gain and loss, respectively) between RNA expression and CNA reported by CHASM (**Fig. 3E, Supplemental Fig. 5C**). For epiAneufinder, there is a significant negative correlation (r = - 0.2) between RNA expression and the extent of CNA loss reported, while the correlation with CNA gain is minimal (**Fig. 3F, Supplemental Fig. 5D**). Since CopyscAT only reports CNA by chromosome, we examine the distribution of scaled chromosomal transcript by copy number state assignment. Indeed, there are significant stepwise increases in gene expression from whole-chromosome deletion to one arm deletion, to one arm gain, and whole-chromosome gain; however, the difference is not evident (**Fig. 3G**).

In addition, Wilson et al suggest that LOY should be a biomarker for genome instability^27^, and thus we hypothesize that cells with a LOY genotype likely contain more CNA. Indeed, CHASM reports a significant higher fraction of genome with CNA detected in the LOY cells compared to the XY genotype (**Fig. 3H**). In both epiAneufinder and CopyscAT, no significant difference is seen between the LOY and XY genotype (**Fig. 3I-J**). Note that the scale of fraction of genome with CNA reported in epiAneufinder and CopyscAT is much higher than CHASM, possibly suggesting the potential false positive detections overwhelm the genuine signals.

Because p53 plays a central role in DNA damage repair, we next compare CNA burden in p53 wild-type and knock-out conditions from multiple tissue types in wild-type (4.5-6 months old) and germline p53 knock-out (4.5–8 months old) mice profiled with Microwell single-cell ATAC sequencing technology^32,33^ (**Fig. 3K**). After per-cell quality control and filtering for tissues with sufficient cell number in both conditions, we focused on thymus, pancreas, and liver (**Supplemental Fig. 6A-C**). In wild-type mice, in which CNA burden is expected to be extremely low because p53 is intact and the animals are young, CHASM detects significantly fewer CNAs (median % genome altered 0-3.52% in CHASM) than the other methods, whose estimated % genome altered is >5% in most tissues (**Fig. 3L**). We next hypothesize that p53 knockout would be associated with higher CNA burden. Indeed, CHASM identifies a pronounced increase in the fraction of cells with CNA in >5% genome in T and B cells, a pattern that is also observed from epiAneufinder results (**Fig. 3M**). In thymus antigen presenting cells (APC) and myeloid cells, however, only CHASM reveals higher CNA burden in p53 knock out mice, whereas the other tools do not (**Fig. 3M**). Intriguingly, both CHASM and epiAneufinder show that hepatocytes from the liver tissue have lower fraction of cells with >5% genome altered in p53 knock out mice, and that no difference is detected in pancreas endocrine cells. Given two of three p53 knockout mice developed lymphoma, it suggests that the highly proliferative cell types like lymphocytes and epithelial cells may be most susceptible to genome instability whereas low proliferative cell types like endocrine cells are less susceptible. Taken together, these results show that CHASM gives the expected low genome instability rate for young wildtype mice and recovers, with high cell type specificity, the expected differences in CNA burden between wildtype and p53 knockout mice.

In summary, here we validate CHASM through cross-modality comparisons and biologically grounded contrasts. Across settings, CHASM consistently recovers CNA signals from scATAC-seq that align with orthogonal measurements and biological expectations, including concordance with matched scDNA and scRNA data, enrichment of genome instability in LOY cells, and increased CNA burden in the context of p53 knockout. These results establish that CHASM can be applied to a wide range of data for reliable CNA detection.

### Age-associated CNA burden patterns across tissues and cell type compartments

Having validated CHASM using controlled spike-ins, matched modalities, and genome-instability contrasts such as p53 deficiency and loss of chromosome Y, we next asked whether CHASM could reveal age-associated CNA burden across tissues. Large-scale studies have established that mCAs increase with age in blood and can arise across normal human tissues, but these analyses have largely relied on bulk data and are therefore powered mainly for clonally expanded events^1,3,11–14^. As a result, the cell-type specificity of age-associated chromosomal mosaicism remains poorly understood. We therefore applied CHASM across multiple aging datasets, using these analyses both to assess whether CNA calls recover coherent aging-associated patterns and to identify the cellular compartments most susceptible to chromosomal alteration with age.

We first examined young and aged mouse hematopoietic stem cells (HSCs)^34^, motivated by extensive human studies showing that hematopoietic mCAs increase markedly with age^7,11–14^. However, these bulk-data studies primarily capture expanded clones and do not resolve CNA burden directly in individual HSCs. CHASM reveals elevated CNA burden across the genome in aged HSCs, including more frequent chromosome X alterations and losses of chromosome 11 and 18 (**Fig. 4A**). Notably, tumor suppressors Trp53 and Pten reside on chromosome 11 and 18, respectively, supporting that the loss of those chromosomes may confer survival benefits. In terms of clonal frequency of cells with high CNA burden, CHASM detects a 2.47-fold increase in the fraction of cells harboring CNAs across more than 5% of the genome in aged HSCs, whereas epiAneufinder detects a 0.95-fold decease and CopyscAT detects no difference (**Fig. 4B, Supplemental Fig 7A**). These results suggest CHASM reveals more plausible aging trends in the HSC compartment.

**Figure 4.**
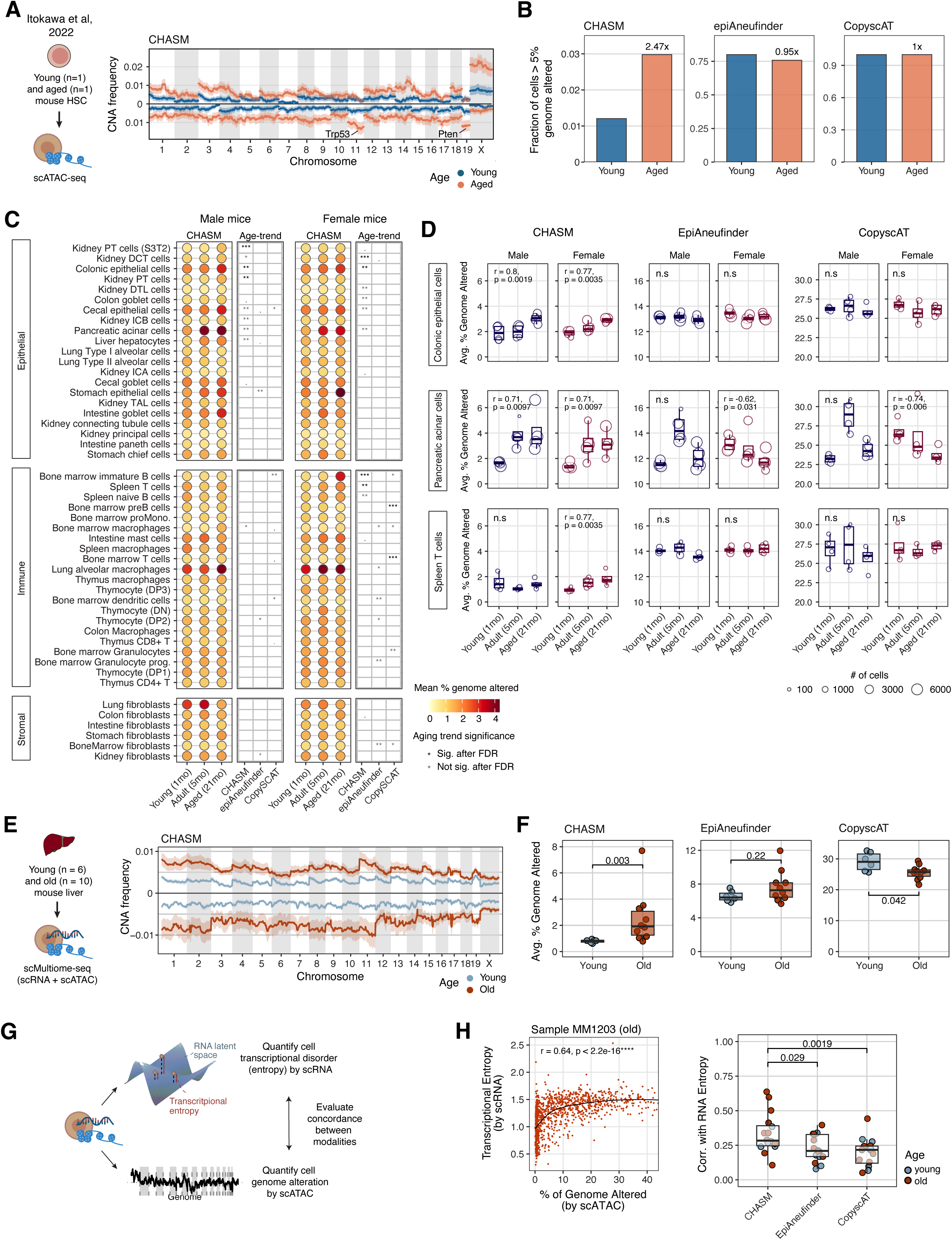
Age-associated CNA patterns across tissues. **A)** Schematic of young and aged hematopoietic stem cell data and CNA frequency from CHASM along the genome. Color indicates age group. Shaded lines represent standard error of CNA frequency across cells. **B)** Bar plots of the fraction of cells with >5% genome altered determined from the results from CHASM, epiAneufinder, and CopyscAT. Fold changes in cell fractions between aged and young samples are annotated. P-values are derived from Chi-squared test. **C)** Bubble plot of the average percentage of genome with CNA in each cell type from samples in the same age group determined by CHASM (left). Cell types with sufficient cells and samples per age group passing quality control are shown. Statistical significance of age-associated increase in CNA burden in each cell type is determined by Spearman correlation (right). Text color indicates the corresponding significance level remains significant after multiple-testing correction. Results on CHASM, epiAneufinder, and CopyscAT are shown. **D)** Boxplots of sample mean percentage of genome altered in colonic epithelial cells, pancreatic acinar cells, and spleen T cells. Each dot is a sample. Dot size indicates the number of cells in each sample. Spearman correlation coefficient and p-values are annotated. **E)** Schematic of scMultiome-seq performed on liver tissue from young and old mice and genome-wide CNA frequency detected in hepatocytes by CHASM. **F)** Box plots comparing the average percentage of genome with CNA determined by CHASM, epiAneufinder, and CopyscAT between young and old mice. Each dot is a mouse. Two-sided Wilcoxon tests are performed. **G)** Schematic of cross-modality comparison between transcriptional entropy estimated from scRNA-seq and the percentage of genome with CNA from scATAC-seq. **H)** Scatter plot of the percentage of genome with CNA by scATAC from each cell detected by CHASM and transcriptional entropy of the same cell by scRNA in an old mouse MM1203. A loess curve is fitted, and the Spearman correlation coefficient is annotated. Box plots comparing the Spearman correlation coefficients between scRNA-estimated transcriptional entropy and scATAC-estimated genome alteration from CHASM, epiAneufinder, and CopyscAT. Two-sided Wilcoxon tests are performed between each of epiAneufinder and CopyscAT against correlation coefficients from CHASM results. Color indicates mouse age group.

We next extended this analysis to a whole-organism mouse aging scATAC atlas spanning young (1 month), adult (5 months), and old (21 months) mice across multiple tissues ^26^. Although aging is generally expected to be associated with increased CNA burden, these patterns have not been characterized at cell-type resolution. Further, since this data was generated via EasySci, a different scATAC-seq protocol^35^, this analysis serves as an additional robustness test across tools. After per-cell quality control and constraining to lineages with sufficient cells and samples, we focus on several visceral organs for the epithelial compartment, immune organs for immune cells, and ones with sufficient fibroblasts for the stromal cells (**Supplemental Fig. 8A**). Overall, CHASM detects low baseline CNA burden across age groups, whereas both epiAneufinder and CopyscAT report more substantial mCA even in young mice (**Fig. 4C-D, Supplemental Fig. S8B-C**). Across methods, monotonic age-associated increases in CNA burden are observed in several cell types such as cecum epithelial cells and lung alveolar cells, but these trends are more frequently detected in CHASM than with the other tools (**Fig. 4C**). After multiple-testing correction, only a subset of these trends remain significant, likely owing to limited sample number.

Broadly, CHASM identifies age-related increase in CNA burden in a subset of tissues and cell types, with signals more frequently observed in epithelial cell types. Specifically, epithelial cells from the kidney, colon and cecum, as well as pancreatic acinar cells, show evident age-associated increase in CNA burden across age groups in CHASM (**Fig. 4C-D**). However, because epithelial compartments often contain larger numbers of cells, this pattern may partly reflect increased detection power. In comparison, the age-dependent trend in CNA burden in epithelial cells are less apparent with epiAneufinder and CopyscAT (**Fig. 4C-D, Supplemental Fig. S8B-C**). In the immune compartment, several cell populations, such as T and B cells in spleen, exhibit more evident age-related CNA burden pattern in females than males (**Fig. 4C-D**). This pattern may relate to observations from Lu et al describing multiple sex- and age-dependent changes in immune cell types, including a female-biased decrease in splenic naïve B cells with age^26^. In summary, CHASM detects age-associated CNA burden increases in several tissue and cell type compartments, including epithelial populations where signals are often observed in both sexes.

We then focused on aging liver, a tissue with distinctive ploidy biology and prior evidence of age- and injury-associated somatic genetic change^36,37^. Because hepatocyte aneuploidy has been debated and whole-chromosome alterations appear rare by single-cell DNA sequencing^38^, this dataset provides a stringent setting to test whether CHASM can detect subtle, age-associated CNA burden while linking it to transcriptional state. Our analysis encompassed hepatocytes from the livers of 6 young and 10 old mice (**Fig. 4E**). CHASM shows significantly higher mean percentage of genome with CNA in old mice compared to young mice, whereas epiAneufinder shows no difference and CopyscAT shows a reversed trend (**Fig. 4F, Supplemental Fig. S7B**). At the level of gene expression, Yang et al. recently linked increased transcriptional noise to aging^39^. We thus test if there is an association between transcriptional noise and genome instability, with the former providing orthogonal evidence of underlying cellular dysregulation associated with aging (**Fig. 4G**). Across individual mice, we see moderate but significant single cell-level correlation between genome instability, as quantified by fraction of genome with CNA detected via CHASM on the ATAC modality, and transcriptional entropy, quantified on the RNA modality (**Fig. 4H**). Notably, CHASM’s estimate of the fraction of genome with CNA shows significantly higher correlation with transcriptional entropy than the estimates produced by epiAneufinder and CopyscAT. This improved correlation suggests that CHASM’s high specificity improves the recovery of biologically meaningful trends.

To summarize, we apply CHASM to quantify age-associated CNA burden across cell types and tissues using data generated by different ATAC-seq technologies. CHASM reveals monotonic age-associated, cell type-specific increases in CNA burden across multiple cell types, whereas these trends are often less evident or absent with other tools. In liver hepatocytes, increased CNA burden is further associated with increased transcriptional disorder. Notably, these patterns coherently emerge only with CHASM, demonstrating that stringent control of false discovery is essential for uncovering biologically meaningful CNA signals in non-neoplastic tissues. These multi-axis validations establish CHASM as a reliable framework for studying chromosomal instability at single-cell resolution.

### Aging kidney tissues harbor cell type and genomic region-specific aneuploidy

The kidney is a well-suited model for studying somatic mosaicism, as it comprises diverse epithelial cell types with varying susceptibility to injury and regenerative capacity, owing to differential exposure to metabolic waste and toxins during urine processing^40^. We curate a kidney scATAC atlas composed of 99 individuals across ages (20-99 years old) and kidney disease conditions (AKI, CKD, and non-disease), 66 of which have cell-matched RNA data available^27,40–44^ (**Fig. 5A**). After batch integration, we annotated cell type using lineage-specific markers and compared our annotations to an existing atlas from the Kidney Precision Medicine Project (**Methods**). We identify nine epithelial cell types (**Supplemental Fig. 9A**), and consistent with previous publications^27,40,45^, there is a cluster of injured proximal tubule cells that express *VCAM1* alongside other injury-associated markers (**Supplemental Fig. 9B**).

**Figure 5.**
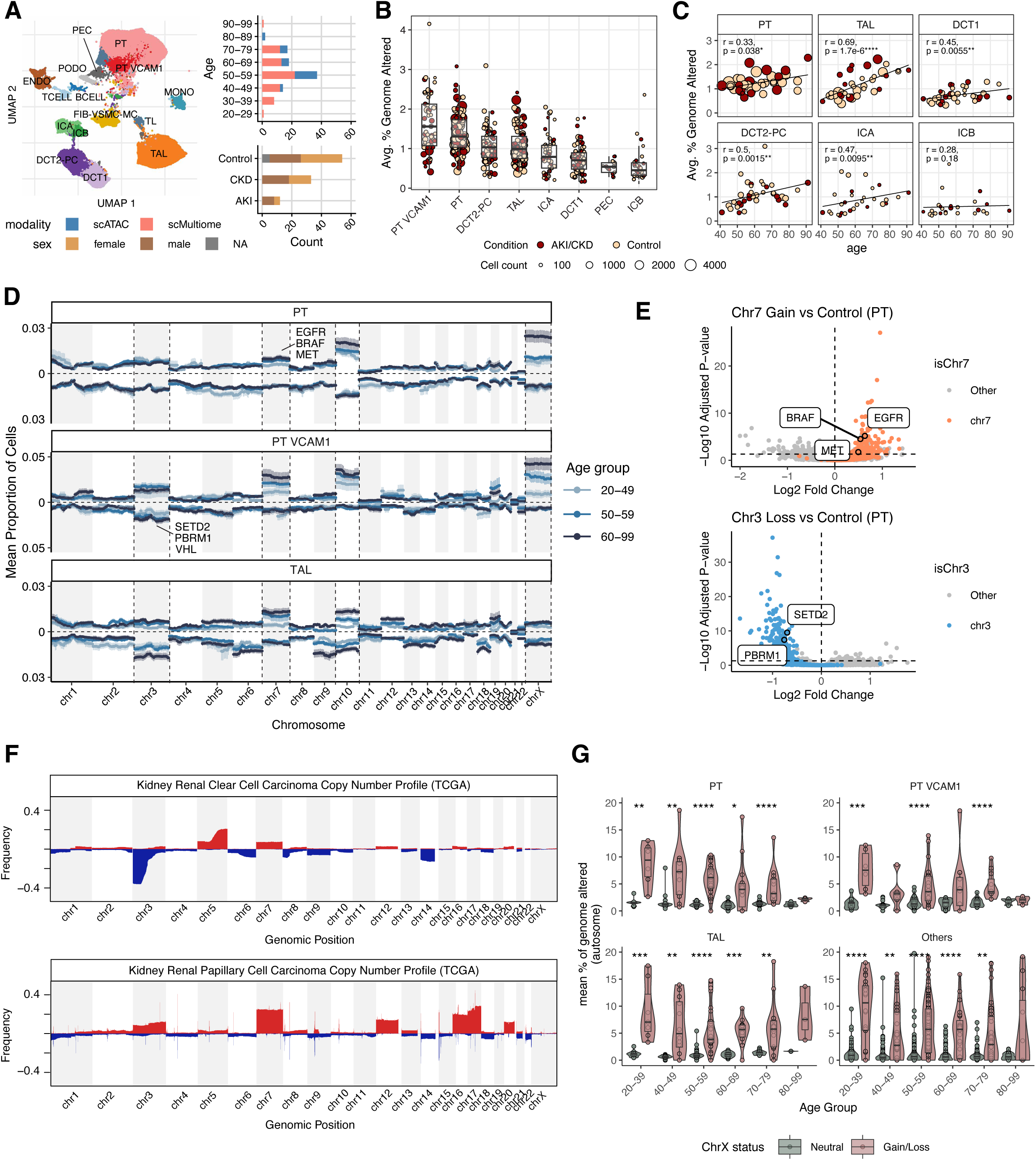
Cell type specific mosaic chromosomal alterations in aging kidney tissues. **A)** UMAP visualization of integrated kidney atlas data. The top-right bar plot shows the number of samples across different age group, colored by the data modality. The bottom-right bar plot shows the number of samples across disease conditions, colored by sex. PCT=proximal tubule, PT_VCAM1=VCAM1+ injured proximal tubule, PEC=parietal epithelial cells, TL=thin limb, TAL=thick ascending limb, DCT1=early distal convoluted tubule, DCT2_PC=late distal convoluted tubule and principal cells, ICA=type A intercalated cells, ICB=type B intercalated cells, ENDO=endothelial cells, PODO=podocytes, TCELL=T cells, BCELL=B cells, MONO=mononuclear cells, FIB_VSMC_MC=fibroblasts, vascular smooth muscle cells and mesangial cells. **B)** Box plots of the average percentage of genome with copy number alteration (CNA) in epithelial cell types. Each dot is an individual. Dot size represents the number of cells in the cell type. Color represents disease condition. **C)** Scatter plots of the average percentage of genome with CNA and individual age, faceted by epithelial cell types. Dot size represents the number of cells in the cell type. Color represents disease condition. **D)** Frequency of cells with CNA at each genomic bin in proximal tubule cells (PT), injured proximal tubule cells (PT VCAM1), and thick ascending limb cells (TAL). The positive direction summarizes copy number gain, and the negative direction summarizes copy number loss. Color indicates age group. The error bars represent standard error in CNA frequency across individuals. **E)** Volcano plots of differentially expressed genes between PT cells with chromosome 7 gain and copy neutral (top), and between PT cells with chromosome 3 loss and copy neutral (bottom). Color indicates whether the gene is in the affected chromosome. **F)** Frequency of cells with CNA at each genomic bin in clear cell renal cell carcinoma (ccRCC) (top) and papillary renal cell carcinoma (pRCC) (bottom) from TCGA. The positive direction summarizes copy gains and colored in red, and the negative direction summarizes copy losses and colored in blue. **G)** Violin plots of the average percentage of the autosomal genome with CNA between cells with and without CNA at chromosome X in each age group. Results on PT, PT VCAM1, TAL, and the rest of the epithelial cell types are shown. Two-sided Wilcoxon tests are performed.

To our knowledge, this represents the first cell-type-resolved characterization of mCA distribution across a large cohort of kidney tissues. CHASM shows that, across cell types, VCAM1-high proximal tubule cells (PT VCAM1) harbor the highest CNA burden, followed by proximal tubule cells (PT), late distal convoluted tubule subtype and principal cells (DCT2-PC), and thick ascending limb (TAL) (**Fig. 5B**). Notably, the non-injury-associated cell types PT, TAL, DCT1, and DCT2-PC show positive correlation between genome-wide CNA burden and age (**Fig. 5C**), consistent with increased genome instability as a hallmark of aging^46^. Here, the PT cells do not include PT VCAM1 cells, and thus the increased CNA incidence with age is not the result of increased presence of injured PT cells. Additionally, age information is not used in the detection of CNAs, thus the recovery of a consistent age-associated trend provides independent biological validation of the signal. In contrast, if the detected events were dominated by noise or technical artifacts, such structured and monotonic associations with age would be unlikely to emerge.

We next examine cell type-specific CNA frequency along the genome, stratified by age group (**Methods**). We divide individuals into three age groups by balancing sample count in each group: 20-49, 50-59, and 60-99 years. First focusing on PT cells, we find a monotonic increase in CNA frequency with age in chromosomes 10 and X, and to a lesser extent in chromosome 7 (**Fig. 5D**). Chromosome 7 harbors several oncogenes such as *EGFR*, *BRAF*, and *MET*, which concordantly show increased expression in PT cells with chromosome 7 gain (**Fig. 5E**). Both gain and loss are observed in chromosome 10 which harbors tumor suppressor *PTEN*, and, indeed, PT cells with chromosome 10 loss have reduced *PTEN* expression (**Supplemental Fig. 9C**). Similarly, chromosome 3 alterations, which are prevalent in kidney cancers and harbor tumor suppressors *SETD2*, *PBRM1*, and *VHL*, show consistent transcriptional effects. Although chromosome 3 loss does not seem to be increased in PT cells, differential gene expression analysis suggests that PT cells harboring chromosome 3 loss concordantly show reduced expression of chromosome 3 genes including these tumor suppressors (**Fig. 5E**).

We next consider the disease-associated PT-VCAM1 cells. Similar to PT cells, PT VCAM1 cells have elevated alterations in chromosomes 10, and X, but show substantially increased frequencies of chromosome 7 gain and chromosome 3 alterations. These trends are consistent across all three age groups and show monotonicity with age (**Fig. 5D**). Notably, PT VCAM1 cells display age associated increase in both gain and loss of chromosome 3. Examining CNA profile in clear cell renal cell carcinoma (ccRCC) and papillary renal cell carcinoma (pRCC) from TCGA^47^ reveals that chromosome 3 loss is enriched in ccRCC whereas chromosome 3 gain is enriched in pRCC, suggesting that both chromosome 3 gain and loss have an important role in kidney cancers (**Fig. 5F**). The observation of chromosome 3 alterations and chromosome 7 gain in PT cells and their increasing incidence in the injured, but non-dysplastic PT VCAM1 population suggest that these alterations may represent early cancer-relevant genomic events prior to malignant transformation. However, not all CNAs in cancer tissues are recapitulated in normal tissues, suggesting that they may be later events closer to or after malignant transformation.

TAL cells also harbor similar CNA as PT cells, including chromosome 3 loss and alterations in chromosomes 7 and 10, with frequencies that increase monotonically with age (**Fig. 5D**). The thick ascending limb plays a critical role in salt reabsorption and is highly metabolically active, making it susceptible to physiological stress and injury in kidney disease^40^. The presence of reproducible, age-associated aneuploidies in TAL cells suggests that this compartment is also vulnerable to chromosomal instability during aging. Although we did not identify a distinct cluster of injured TAL in our dataset, prior studies have reported a maladaptive TAL cluster with a transcriptional profile that mimics injured PT^40^. In contrast, DCT1, DCT2 and PC exhibit distinct CNA profiles with higher prevalence of chromosomes 14 and 18 loss (**Supplemental Fig. 9D**). Furthermore, these CNAs tend to occur in different cells rather than co-occurring in the same cell, as most aneuploid cells only harbor CNA in one chromosome, suggesting independent origins of these events (**Supplemental Fig. 9E**). Despite these differences, certain alterations, including those affecting chromosomes 7 and 10, are shared across multiple epithelial cell types, pointing to common axes of genome instability in aging kidney tissue (**Supplemental Fig. 9F**).

Given both PT and PT VCAM1 cells show evident alteration in chromosome X, we next ask whether chromosome X alteration may be associated with broader somatic alteration. Indeed, cells with chromosome X alteration show higher extent of average autosomal mCA burden than cells from the same individual without chromosome X alteration (**Fig. 5G**). This trend holds true across age and epithelial cell types. However, examination of gene expression between cells with and without chromosome X alteration reveals few enriched pathways (**Supplemental Fig. 9G**). Likewise, in cancer patients, individuals with X chromosome alterations are likely to harbor autosomal mosaic events^48^, supporting an association between mosaic X-chromosomal dysregulation and somatic alterations.

In summary, we resolve, at single-cell resolution, the cell type–specific landscape of mCAs in non-cancer kidney epithelium. A small subset of cells within specific epithelial compartments exhibits recurrent mCAs that increase in frequency with age and localize to distinct genomic regions. These alterations are accompanied by concordant transcriptional changes and overlap with regions commonly altered in kidney cancer, suggesting their potential origin in normal tissues. Additionally, chromosome X alteration may emerge as a potential biomarker of somatic alterations.

### Elevated CNA burden in non-disease cells is associated with features of injury

We next characterize, at single cell resolution, transcriptomic and chromatin features associated with increased CNA burden to investigate the biological states of these altered cell populations. We define CNA burden as the proportion of the genome with CNAs. Visualizing CNA burden on the weighted nearest neighbor-based UMAP does not reveal distinct cell clusters (**Fig. 6A**), suggesting the effect of CNA on cellular phenotype may be heterogeneous and subtle. To investigate genes associated with CNA, we correlate CNA burden in each cell, computed from the ATAC-modality, with gene expression in the matched RNA modality, stratified by cell type and individual. The individual-level correlations are then combined across individuals through a meta-analysis (**Methods**). Given that PT and TAL cells are the two most abundant cell types in the kidney epithelium, our analyses are focused on these two cell types. In PT cells, healthy PT marker genes such as *MME*, *HNF4A*, and *CUBN* show negative correlation with CNA burden, whereas injury marker genes such as *HAVCR1, VCAM1, ADAMTS1*, and *JUNB* show positive correlation (**Fig. 6B**). Additionally, *TP53* and *CDKN1A* (p21) also show positive correlation with mCA burden in PT cells, suggesting a role of DNA repair response and senescence (**Supplemental Fig. 10A**). Similarly, in TAL cells, healthy TAL markers such as *ESRRG, EGF* are negatively associated with CNA burden, whereas *VCAM1, CD24, S100A6*, which are markers for adaptive and degenerative TALs^49^, are positively correlated with CNA burden (**Fig. 6B**). Pathway enrichment analysis reveals TNF*α* signaling via NF*κ*B, epithelial to mesenchymal transition (EMT), interferon gamma and alpha response, apoptosis pathways are significantly enriched in CNA-associated genes (**Supplemental Fig. 10A**).

**Figure 6.**
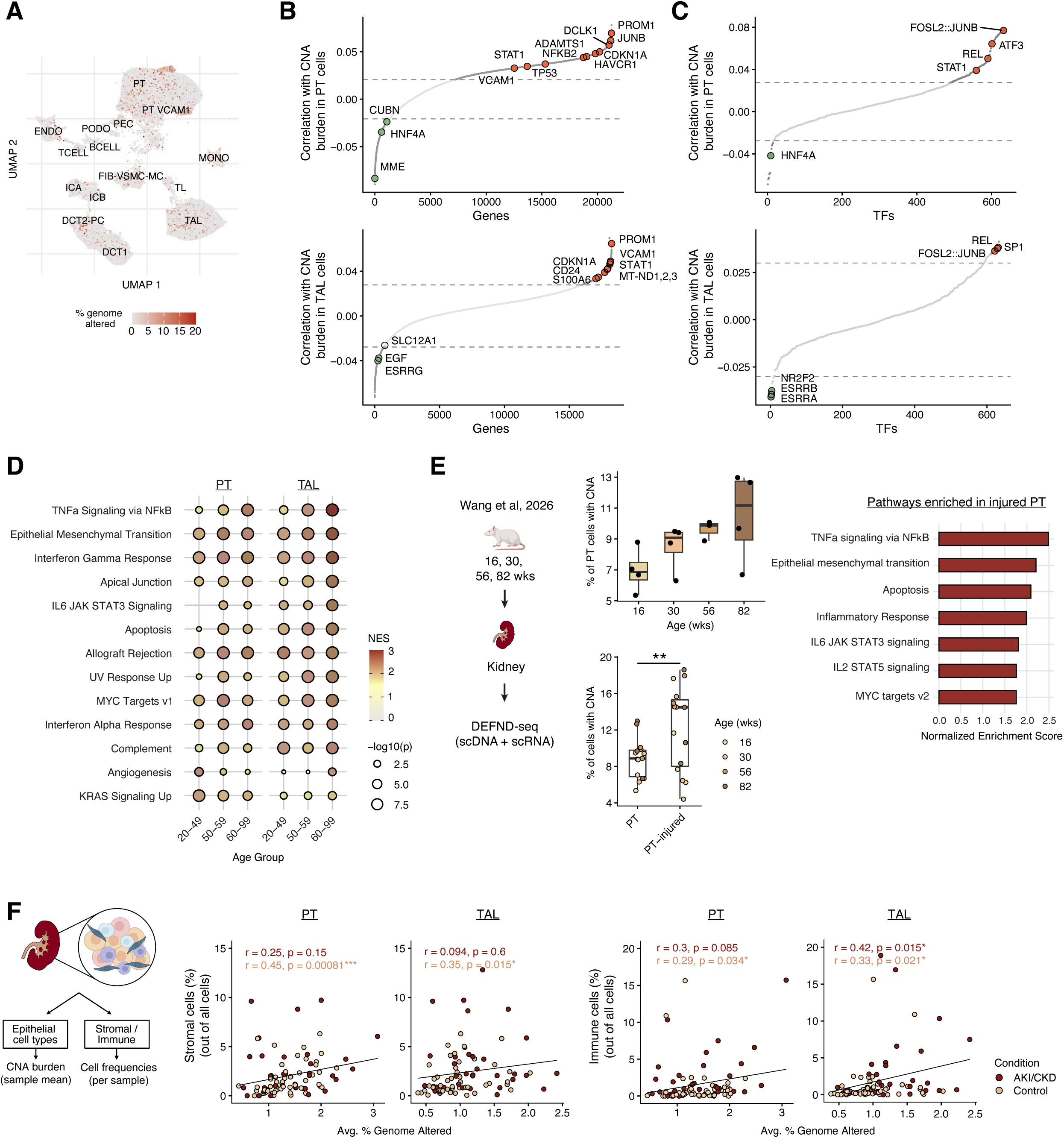
Cellular and tissue features associated with mosaic chromosomal alteration burden. **A)** UMAP of integrated kidney atlas colored by the percentage of genome with copy number alteration (CNA). **B-C)** Rank plot of gene correlation coefficient between CNA burden in cells and the gene’s expression level **(B)** and transcription factor activity **(C)**. Top panels show results on proximal tubule cells (PT) and bottom panels shows results on thick ascending limb cells (TAL). Orange dots indicate injury marker genes in each cell type. Green dots indicate marker genes in healthy cells. Dotted lines show the significance thresholds. **D)** Pathway enrichment analysis on genes correlated with CNA burden in PT and TAL. Color indicates net enrichment score. Dot size indicates p-value. **E)** Schematic of DEFND-seq data on aging rat kidney tissues^53^. Top box plot shows the percentage of cells with CNA detected from DEFND-seq in each age group. Bottom box plot shows the percentage of PT and injured PT cells with CNA detected from DEFEND-seq. Two-sided Wilcoxon test is performed. Pathway enrichment analysis result on genes more highly expressed in injured PT cells (right bar plot). **F)** Schematic of data analysis workflow. The percentage of genome with CNA (CNA burden) is averaged within each sample and epithelial cell type, and the frequencies of stromal and immune cells are calculated. Scatter plots between CNA burden and the frequency of stromal cells (left) and immune cells (right). Each dot is a sample. Color represents disease condition. Spearman correlation coefficients are calculated per disease condition.

To ensure the robustness of these results, we perform the same analysis independently within each age group and evaluate replicability across age groups. Among these, TNF*α* signaling via NF*κ*B hallmark pathway is consistently enriched in CNA-elevated genes in both PT and TAL cells, and the enrichment score further increases monotonically with age (**Fig. 6D**). Similarly, EMT, IFN*λ* and IFN*α* response, apoptosis pathways remain the top pathways enriched for association with mCA across age and cell types. The consistently replicable enrichment of IFN response and EMT pathway suggest that cells harboring higher CNA burden correspond to heightened transcriptional features of inflammation and increased adaptive plasticity.

We next examine transcription factors associated with CNA. We utilize chromVAR^50^ to compute a TF binding score across the genome for each cell and perform correlation analysis across cells (**Methods**). This reveals that the accessibility at binding sites of AP1 family transcription factors (e.g. JUN/FOS) and REL (a member of the NF-*κ*B family) are positively correlated with mCA burden in PT and TAL cells (**Fig. 6C**). These TFs are well known mediators of stress, inflammatory, and injury responses and have been shown to be more active in injured PT and injured / adaptive TAL^40^. In contrast, HNF4A, a key regulator of proximal tubule differentiation and metabolic function^51^, shows a negative correlation with CNA burden in PT cells, while ESRRB, which are involved in mitochondrial metabolism and maintenance of epithelial function^52^, are similarly negatively correlated with CNA in TAL cells. Together, these patterns suggest that increased CNA burden is associated with activation of injury- and inflammation-related regulatory programs and a concomitant loss of differentiated epithelial cell identity.

To investigate if the CNA-related transcriptional features identified in the human atlas data reflect conserved biology, we turn to an aged rat model and performed DEFND-seq (scDNA and scRNA co-assay) on kidney tissue^53^. In this data, non-injured PT cells show increased CNA burden with age (**Fig. 5E**). Injured PT cells have a higher burden of chromosomal CNA compared to non-injured PT cells, consistent with our results in human samples (**Fig. 6E**, **Fig. 5C**). Furthermore, these injured kidney cells have higher expression of injury markers including *Adamts1*, *Havcr1*, and lower expression of healthy markers *Hnf4a*, *Mme*, and *Cubn* (**Supplemental Fig. 10B**), consistent with genes showing higher association with CNA burden (**Fig. 5B**). Interestingly, *Prom1* is more highly expressed in non-injured PT cells, yet it shows a positive correlation with CNA burden in the human samples. Given its role in tissue regeneration, this may suggest that PT cells with higher CNA burden are engaged in processes of repair and adaptation. At the pathway level, pathways upregulated in injured PT cells in rat coincide with pathways correlated with high CNA burden in human data. The convergence of CNA associated pathways and injury features in both human and rat suggest the phenomenon is biologically conserved. Interestingly, *DCLK1*, associated with CNA burden in human data (**Fig. 6B**), is upregulated in injured PT cells in rat (**Fig. 6E)**. In a companion manuscript^53^, we show that overexpression of *Dclk1* induces higher expression of inflammatory genes such as TNFa and stress response markers such as *Junb* and *Atf3*^53^. *DCLK1* inhibitors have been investigated as possible anti-inflammatory agents in acute lung injury^54^ and may be a targetable pathway in kidney disease.

To investigate whether mCAs in PT and TAL cells may be associated with an altered tissue microenvironment, we characterize tissue cell type composition in relation to their CNA burden. This reveals that in individuals without kidney disease, higher average CNA burden in PT and TAL cells are positively associated with higher proportion of stromal and immune cells (**Fig. 6F**). Consistent with increased presence of stromal cells, PT and TAL cells with high CNA burden show higher expression of fibrotic markers, suggesting that these genome-altered epithelial cell populations engage in fibrosis that may be related to maladaptation (**Supplemental Figure 10C**). Additionally, among individuals without kidney disease, higher CNA burden in PT cells is correlated with increased presence of myeloid cells, whereas CNA burden in TAL cells tend to have higher T cell presence (**Supplemental Figure 10D**).

Overall, at the single cell level, epithelial cells with higher CNA burden exhibit moderate but consistent increases in transcription of injury markers, inflammatory response, and stress-associated TF activity, even when they do not cluster with overtly disease-associated cell types. At the tissue level, in individuals without clinical disease, higher average CNA burden within PT and TAL cells is associated with immune and stromal infiltration. Together, these findings suggest that mCAs in specific epithelial subsets are linked to early, pre-disease cellular states and may contribute to a more fibrotic and inflamed tissue microenvironment, with potential relevance to the initiation of kidney disease.

## DISCUSSION

With growing interest in somatic mosaicism and cellular aging, along with large-scale atlas efforts generating single-cell data, there is a critical need for methods that can detect somatic mutations while simultaneously resolving cell identity and state. Understanding which cell types and cellular states harbor genomic alterations is essential for elucidating their functional consequences in aging tissues. However, bulk sequencing lacks cell type resolution, and single-cell DNA sequencing, while precise, is difficult to scale and does not directly capture cell state. In contrast, single-cell ATAC-seq provides a scalable modality that reflects cell identity and regulatory state and can be integrated with gene expression through multiome assays. Here, we develop a framework for high-specificity detection of CNA signals from scATAC-seq data, applicable to non-dysplastic tissues where CNAs are non-clonal and thus low-frequency. We show that existing methods for CNA detection in scATAC-seq data exhibit high false discovery rates, largely due to insufficient modeling of background variations that can be misinterpreted as CNA signal. Such high false discovery rates can overwhelm and obscure low-frequency CNA signals in normal aging tissues.

The core idea of CHASM is to estimate a cell-matched normal baseline, rather than relying on the identification of a null population or a cross-population average baseline. Because chromatin accessibility varies across cell states and genomic regions, heterogeneous sources of variation must be accounted for when constructing an appropriate control. To this end, we use wavelet transformation coupled with robust PCA to separate background variation from real CNA events. By transforming the data into a representation where CNA signals are sparse and then isolating them from background variation, CHASM improves the accuracy of CNA detection in heterogeneous single-cell data. Extensive benchmarking using spike-in experiments, cross-modality comparisons, and comparisons between cells with expected differences in genome instability demonstrates that CHASM preserves sensitivity while effectively reducing false positives, enabling confident analysis of normal tissues where CNAs are rare.

Beyond benchmarking, CHASM enables a cell type–resolved view of how chromosomal instability accumulates during aging. Prior studies have established that mCAs increase with age in blood and across bulk tissues^1,3,11–14^, but these approaches cannot identify the cellular compartments in which alterations arise. Applying CHASM across aging datasets, including hematopoietic stem cells, a whole-organism mouse aging atlas, and liver multiome data, reveals that age-associated CNA burden is not uniform across the genome nor across cell lineages. Instead, specific cell populations accumulate CNAs with age, whereas others appear comparatively resistant. In hepatocytes, increased CNA burden is linked to transcriptional disorder measured independently from the RNA modality. These analyses extend prior bulk studies by resolving the cell-type landscape of genome instability during aging and establish aging-associated chromosomal mosaicism as a heterogeneous, cell-context-dependent process.

Having established that CHASM recovers coherent age-associated CNA patterns across tissues, we next used human kidney as a disease-relevant setting to define how chromosomal mosaicism manifests within an aging epithelial organ. Across 99 human kidney samples spanning the lifespan and kidney disease status, CHASM reveals a non-random, cell type–specific landscape of rare chromosomal alterations. These events increase with age, preferentially affect specific epithelial compartments, and localize to genomic regions recurrently altered in kidney cancer. A previous pan-tissue study that inferred mCA based on bulk RNA-seq data struggled to detect such events in kidney tissue^3^, likely due to their low frequency. In contrast, CHASM applied to scATAC-seq data enables detection of these rare events and their assignment to specific cell types.

The presence of cancer-associated CNAs in non-diseased tissues may point to early alterations that may contribute to increased risk as well as potential precursors to clonal expansion. We observe that injured PT cells harbor higher frequencies of CNA on chromosomes 3 and 7, alterations commonly observed in kidney cancers, and that carrier cells exhibit corresponding transcriptional changes in the cancer genes located within these regions as compared to non-carrier cells. PT cells are thought to be the cell of origin for both clear cell renal cell carcinoma^55^ and papillary renal cell carcinoma^56^, and injured PT cells have been shown to adopt a more dedifferentiated and proliferative state^57^. Together, these findings suggest that aging tissues provide a window into early cellular changes preceding overt disease, capturing molecular alterations that may seed malignancy. Characterizing such cell populations improves our understanding of early disease processes and the transition from normal aging to disease.

We show that PT and TAL cells with increased mCA burden exhibit elevated inflammatory response, with the NF *κ* B response pathway among the most strongly enriched programs associated with mCA burden. This is consistent with a broad link between genome instability and inflammation^58^. An increased mCA burden may reflect a higher level of genome instability, possibly leading to accumulation of cytoplastic DNA that is sensed by cGAS, activating STING and downstream NF*κ*B signaling^58^. In addition, DNA damage response pathways can directly activate NF *κ* B signaling. Studies have investigated the role of DNA damage in driving inflammatory responses^58^. The association between aging, DNA damage response, and inflammation is core to the concept of inflammaging, which is a chronic, systemic inflammation that increases with age and contributes to age-related diseases^59^. In contrast to acute inflammation, which facilitates pathogen clearance, chronic inflammation can lead to tissue damage and may engage feedback mechanisms that dampen immune responses, potentially promoting immune evasion rather than enhanced surveillance^60^. While the triggers that initiate mCA remain unclear, their downstream effects may contribute to the establishment of a pro-inflammatory tissue environment that promotes maladaptive cellular states.

As aging is associated with increased mCA burden, it reflects a process beyond transient stress responses and instead involves the accumulation of structural alterations to cellular integrity that may be difficult to reverse. Reducing the impact of chronic inflammation is therefore of potential translational interest. *DCLK1* is among the top genes associated with CNA burden in PT cells. It is a protein kinase which has been shown to participate in DNA damage response pathways and can influence downstream NF-*κ*B signaling^61^. Notably, inhibitors targeting *DCLK1* have been explored for their potential to modulate inflammatory responses in acute lung injury as well as to inhibit tumorigenesis in kidney^62^. Similarly, *STAT1* expression is also associated with mCA burden in both PT and TAL cells. Inhibitors that target JAK/STAT signaling axis have also been investigated for alleviating chronic kidney disease^63^. These findings provide a foundation for designing intervention to target the impact of mCA accumulation and chronic inflammation.

Because CHASM is applied to scATAC-seq, the inferred events reflect large-scale deviations in accessible chromatin. Many of these likely correspond to underlying DNA copy-number alteration; however, broad chromatin remodeling without changes in DNA content could also produce similar signals. Conversely, true DNA CNAs that are buffered by compensatory changes in chromosome accessibility may be under-detected. We therefore interpret CHASM calls as large-scale dosage-like alterations in accessible chromatin, with substantial but not universal correspondence to DNA copy number. Sex chromosomes present an additional challenge. In females, one copy of the X chromosome is typically inactivated and inaccessible^48^, and in males, much of the Y chromosome is relatively heterochromatic^64^. As a result, alterations in sex chromosomes inferred from chromatin accessibility may not consistently reflect underlying DNA-level changes. At the same time, large-scale changes in sex chromosome accessibility remain an active area of investigation^48,65^. Specifically, X chromosome inactivation is essential for normal cellular function in females, and aging may be associated with loss of this inactivation^65^, which could manifest as increased X chromosome accessibility. In this context, integrating both DNA- and ATAC-based measurements will be important for more precise interpretation.

We demonstrate that chromosome X alterations increase with age and are associated with increased autosomal alterations. In a study on cancer patients using SNP arrays, chromosome X alterations occur at a higher rate than autosomal events^48^, similar to our observations in the kidney epithelium. As for the association with autosomal mosaic events, one possible explanation is that sex chromosomes are more permissive to dosage variation due to mechanisms such as X chromosome inactivation, allowing cells harboring such alterations to persist, whereas comparable alterations in autosomes may be less tolerated. In this context, chromosome X alteration may indicate broader genome instability, analogous to previous observations of chromosome Y loss^8,27^.

Large efforts like the Somatic Mosaicism across Human Tissues Network (SMaHT) and Sennet consortium^24,25^, will be critical for developing effective tools to characterize of aging-related somatic mutations across tissues and populations. Integration with spatial profiling will further enable mapping of mosaic alterations within intact tissue architecture, providing insight into how genome instability is distributed across microenvironments and how altered cells interact with their neighbors. The core concept underlying CHASM, which infers cell-specific baselines for precise accounting of background variation, may also extend to these settings, where heterogeneity in cell composition and state is intrinsic and reference populations are not readily defined.

## ACKNOWLEDGMENTS

We would like to acknowledge NIH support to Dr. Parker Wilson (NIH K08DK126847, NIH R03DK144130) and to Dr. Parker Wilson and Dr. Nancy Zhang (NIH R21DK142089), and NIH support to Dr. Nancy Zhang (1R56AG081351, R01GM25301, R01GM149671). This work was also funded in part by a pilot award to Dr. Parker Wilson from the diabetes research center (DRC) at the University of Pennsylvania Institute of Diabetes and Metabolism (NIH DK19525). Marcos G. Teneche was supported by the California Institute for Regenerative Medicine (CIRM) grant EDUC4-12813 and by an American Society of Hematology (ASH) Hematology Inclusion Pathway (HIP) Graduate Student Award. This work was also supported by the NIH Common Fund’s Cellular Senescence Network (SenNet) program to Dr. Peter Adams (grant U54AG079758).

## AUTHOR CONTRIBUTIONS

Conceptualization, P.C.W. and N.R.Z.; Data generation and collection: P.C.W, H.W., M.G.T, and X.E.C.; methods development and implementation: X.E.C. and N.R.Z.; design of computational experiments, X.E.C., and N.R.Z.; data analysis and interpretation, X.E.C., H.W, Y.Y. with feedback from P.D.A, P.C.W, and N.R.Z.; manuscript writing: X.E.C. and N.R.Z., supervision P.C.W., N.R.Z.

## DATA AVAILABILITY

This study analyzes a combination of publicly available datasets and newly generated data. Public datasets used in this study include breast tissue scDNA/scATAC co-assay data from PRJNA1003661 and GSE272504; human kidney single-cell multiome data from GSE232222; p53 wild-type and knockout mouse scATAC-seq data from GSE217661; mouse hematopoietic stem cell scATAC-seq data from GSE190424; processed mouse aging atlas chromatin accessibility data from GSE288730; and renal cancer copy-number alteration data from TCGA accessed through the Broad Institute FireBrowse portal. Additional public human kidney single-cell datasets used to construct the integrated kidney atlas are described in the Methods and corresponding references.

Newly generated data reported in this study will be made publicly available upon publication. Any additional processed data files required to reproduce the analyses will be made available upon publication or from the corresponding authors upon reasonable request.

## SOFTWARE AVAILABILITY

CHASM is available at https://github.com/EmiliaCXY/CHASM. Code used to generate the human kidney atlas is available at https://github.com/GaryWang7/Wang_RAGE24_2026.

## SUPPLEMENTAL FIGURE xxLEGENDS

**Figure S1. Benchmarking of CNA detection performance using spike in copy number alteration in chromosome 1. A-C)** Bar plots of copy number alteration frequency for 1%, 5%, and 10% cells harboring the event. Each row in the panel represents a different copy number alteration event, from two-copy loss to two-copy gain. The red dotted line indicates the expected copy number alteration frequency in chromosome 1. Positive direction summarizes copy number gain. Negative direction summarizes copy number loss. Results from CHASM is shown in A), epiAneufinder in B), and CopyscAT in C).

**Figure S2. Benchmarking of CNA detection performance using spike in copy number alteration in the short arm of chromosome 1 (chr1p). A-C)** Bar plots of copy number alteration frequency for 1%, 5%, and 10% cells harboring the event. Each row in the panel represents a different copy number alteration event, from two-copy loss to two-copy gain. The red dotted line indicates the expected copy number alteration frequency in chromosome 1p. Positive direction summarizes copy number gain. Negative direction summarizes copy number loss. Results from CHASM are shown in A), epiAneufinder in B), and CopyscAT in C).

**Figure S3. Benchmarking of CNA detection performance using spike in copy number alteration in a 50 Mb region in chromosome 1. A-C)** Bar plots of copy number alteration frequency for 1%, 5%, and 10% cells harboring the event. Each row in the panel represents a different copy number alteration event, from two-copy loss to two-copy gain. The red dotted line indicates the expected copy number alteration frequency in the 50Mb region in chromosome 1. Positive direction summarizes copy number gain. Negative direction summarizes copy number loss. Results from CHASM are shown in A), epiAneufinder in B), and CopyscAT in C).

**Figure S4. CNA detection performance on spike-in copy number alterations by parameter sweeping and evaluation on break point detection. A-C)** Pareto frontier of bin-level precision and recall of CNA detection by sweeping a segmentation stringency parameter in CHASM and epiAneufinder. Each row represents different copy number alteration carrier cell frequency (1%, 5%, and 10%). Each column represents different copy number alteration events (from two-copy loss to two-copy gain). Color represents different methods. A) shows assessment on chromosome 1p spike-in. B) shows assessment on 50mb region in chromosome 1. C) shows assessment on chromosome 1. **D-E)** Violin plots on p-value derived from t-tests to compare normalized read count in bins before and after the designed breakpoint in spike-in cells. Each row and column setup is the same as A-C). Two-sided Wilcoxon tests are performed. D) shows result on chromosome 1p spike-in, and E) shows results on 50 mb region in chromosome 1.

**Figure S5. Cross-modality comparisons of CNA detected from scATAC-seq data. A)** UMAP visualization of breast tissue scATAC-seq data colored by cell type marker gene chromatin accessibility. LumSec markers include SAA2 and PI3. LumHR markers include TFF1 and MS4A7. Basal markers include KRT14 and CNN1. **B)** Violin plots of the fraction of genome with discordant copy number state assignment from scATAC-seq and scDNA-seq in each sample. **C)** Density contour plots showing concordance between the fraction of chromosome with copy number alteration inferred from scATAC-seq by CHASM or epiAneufinder and normalized chromosomal transcript count from scRNA-seq in each sample. The highest density regions at 95%, 80%, 60%, 40%, 20%, and 5% are plotted. Red colors indicate copy number gain, and blue colors indicate copy number losses. Spearman correlations are annotated.

**Figure S6. p53 knock-out mouse dataset overview. A)** Violin plots on total fragment per cell from each sample. The dotted line represents the 3,000-fragment threshold used for quality control. **B)** Violin plots on median fragment distance to nearest TSS per cell from each sample. The dotted line represents the 30kb threshold used for quality control. **C)** Stacked bar plots whoing the number of cells that passed quality control per lineage. The dotted line shows the 100 cells threshold used for lineages to perform analysis.

**Figure S7. CNA profile in aged hematopoietic stem cells and liver hepatocytes detected by epiAneufinder and CopyscAT. A-B)** CNA frequency from epiAneufinder (A) and CopyscAT (B) results along the genome from the young and aged hematopoietic stem cell dataset^34^. **C-D)** CNA frequency from epiAneufinder (C) and CopyscAT (D) results along the genome from the aging liver scMultiome dataset.

**Figure S8. Whole-organism aging mouse tissue sample overview and CNA accumulation pattern detected by epiAneufinder and CopyscAT. A)** Stacked bar plots showing the total number of cells that passed quality control in each tissue and cell type. Total cell counts are annotated. Color represent age group. **B-C)** Bubble plot showing average percentage of genome altered computed from epiAneufinder (B) and CopyscAT (C) results in each cell type across samples in the same age group. Aging trend statistical significance is computed by Spearman correlation and is annotated on the right.

**Figure S9. Cell-type specific mosaic chromosomal alterations in aging kidney tissues. A)** Cell type marker gene expression across kidney cell types. Dot size represents the percentage of cells in the given annotation express a given gene. Dot color represents the scaled average expression of the given gene. **B)** Dot plot showing healthy and injured proximal tubule (PT) cell marker gene expression in PT and injured PT cells. **C)** Volcano plots of genes differentially expressed between PT cells with chromosome 10 loss and copy neutral PT cells. **D)** Frequency of cells with copy number alteration at each genomic bin in early distal convoluted tubule cells (DCT1), late distal convoluted tubule cells and principal cells (DCT2-PC), and type A intercalated cells (ICA). The positive direction summarizes copy number gain, and the negative direction summarizes copy number loss. Color indicates age group. **E)** Histograms of the number of chromosomes with CNA incidence across cell types. **F)** Co-occurrence frequency of CNA incidence within the same cell across genome. **G)** Volcano plots of differentially expressed genes between PT cells with chromosome X gain and copy neutral. Bar plot of pathway enrichment analysis on the differentially expressed genes in PT cells with chromosome X gain, excluding the genes on chromosome X.

**Figure S10. Gene expression pathways and tissue features associated with mosaic chromosomal alteration burden in proximal tubule cells and thick ascending limb cells. A)** Gene set enrichment analysis result on genes correlated with CNA burden in PT and TAL cells across the 66 individuals with RNA data available. **B)** Rank plot of correlation coefficient between CNA burden in cells and gene expression, highlighting fibrotic marker genes. **C)** Dot plot showing healthy and injured proximal tubule (PT) cell marker gene expression in PT and injured PT cells from rat kidney tissue. **D)** Heatmap of spearman correlation between CNA burden and percentage of each immune cell type. Scatter plots between CNA burden and percentage of myeloid and T cells are shown for PT and TAL cells in each tissue category.

## METHODS

### Breast tissue scDNA preprocessing and copy number analysis

Breast tissue scDNA/scATAC coassay data were obtained from PRJNA1003661 and GSE272504. Single cell FASTQ data from 18 samples were aligned to hg19 using bowtie2 (v2.5.4)^66^ and duplicate reads were marked using sambamba (v1.0.1)^67^. Paired or single alignment setting was selected based on sample. The copy number analysis strategy was adapted from the original publication^17^ and its accompanying code repository (https://github.com/navinlabcode/normalbreastDNA). Copykit (v0.1.3)^68^ was used with the default bin size of 220kb to summarize read depth across the genome, and cells were required to have reads in at least five bins. Diploid cells were identified using `findAneuploidCells()` without simulation (`remove_XY = F, simul = F`). Single-cell quality metrics were calculated with `runMetrics()` and cells with more than 15 breakpoints were removed. In addition, cells with segment ratios between 0.7 and1.3 across all bins, as well as cells identified as outliers by `findOutliers()` with a resolution of 0.7 were filtered.

### Breast tissue scATAC datasets and preprocessing

The scATAC-seq data processing strategy was adapted from the original publication^17^ and its accompanying code repository (https://github.com/navinlabcode/normalbreastDNA). scATAC data was preprocessed using ArchR (v 1.0.2)^69^ and dimensionality reduction was performed with iterative latent semantic indexing. Batch integration was performed using Harmony. Peaks were called using MACS2 (v 2.2.9.1)^70^. Cell type was annotated based on chromatin accessibility at epithelial lineage marker genes.

### CHASM negative binomial model for chromosomal-level copy number alteration detection

Chromosome-wide total read count was modeled using a negative binomial distribution. Specifically, for a cell *c* and chromosome *r*, the observed read count *X*_*cr*_ was assumed to follow *X*_*cr*_ ∼ *NegBinom*(*α*_*c*_*β*_*r*_*γ*_*cr*_, *ϕ*_*cr*_), where *α*_*c*_ is the (observed) library size of the cell *c*, *β*_*r*_ is the expected fraction of reads mapping to chromosome *r* in the diploid setting, *γ*_*cr*_ is the underlying true copy number for chromosome *r* in cell *c*, and *ϕ*_*cr*_ is the dispersion. We estimate *β*_*r*_ by *β̂*_*r*_, the median fraction of reads mapping to chromosome *r* among the cells of the same type. To estimate dispersion, the null expected read count, 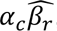, across all cells and chromosomes were partitioned into bins with each bin spanning 5% of the range across all values. The bin size can be adjusted by users. Bins with fewer than 200 values were merged with neighboring bins. For each bin *i*, the dispersion parameter was estimated as 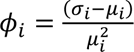 where *μ̂* and *σ_i_* are the’ truncated mean and variance of *X*_*cr*_ whose *α*_*c*_ *β̂*_*r*_ fall into bin *i*. This estimation was done across all cells of the same type. To ensure robust estimation, values in the top and bottom 5% of the read count distribution are excluded. This truncation threshold can also be adjusted by users. To test the null hypothesis of *γ*_*cr*_ = 1 which corresponds to the copy-neutral state, we calculated the probability of observing *X*_*cr*_ under 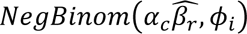. Benjamini-Hochberg procedure was used to control the false discovery rate. Chromosome-level copy number was then estimated as 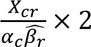.

### CHASM wavelet transformation and robust PCA normalization for sub-chromosomal copy number alteration detection

We first constructed a cell-by-genomic bin count matrix, *X* = [*X*_*cb*_] where *c* indexes cells and *b* indexes genomic bins. Genomic bin size can be specified by users. In our analyses, 2Mb and 5Mb bins were used depending on data quality. 2Mb bins were used in human breast tissue and mouse liver tissue scATAC data. 5Mb bins were used in human kidney tissue scATAC data. Counts were normalized by dividing by the total library size of a cell, applying a square-root transformation, and centering each bin across cells.

To perform wavelet transformation, we constructed a genome-wide, chromosome-arm-specific wavelet basis matrix *W*. For each chromosome arm *r*, a Haar wavelet matrix *W*_*r*_ of size 2^*k_r_*^ was generated using `GenW` from the `wavethresh` package, where 2^*k_r_*^ was the nearest power of 2 greater than or equal to the number of bins in that chromosome arm. Chromosome arms with fewer than 2 bins were excluded from this matrix. In human data, chromosomes 21 and 22 were modelled by full chromosome due to generally lower total read count. Read counts for each chromosome arm were padded with 0 accordingly. These chromosome-arm-specific wavelet basis matrices were stacked diagonally to create the genome-wide chromosome-arm-delineated Haar basis matrix:

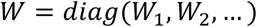

Multiplication XW projected each cell on to this basis and transformed the cell-by-bin matrix into a cell-by-wavelet-basis matrix.

Robust PCA was then performed on of XW using rrpca() from rsvd package, with penalty parameter of

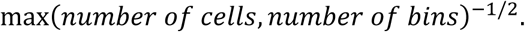

This resulted in two matrices, one representing the core variation and the other representing the signal. Both matrices were transformed back to the linear genome using the inverse wavelet transformation defined by *W*^−1^. DNACopy(v1.78.0)^71^ with significance threshold of 0.005 was then used to segment the read depth residuals derived from the inverse-transformed signal matrix. Users can adjust the significant threshold within DNACopy to get a finer or coarser segmentation.

To assign copy number state, normalized expected and observed read count values were converted back to counts by reversing the centering step, squaring, and multiplying by its library size. For each segment detected by DNACopy, weighted average read counts were calculated for both expected and observed values, using weights corresponding to the contribution of each bin to the total segment depth from the expected values derived from robust PCA. Segment-level copy number state was defined as the ratio of observed to expected weighted average read depth, rounded to the nearest integer:

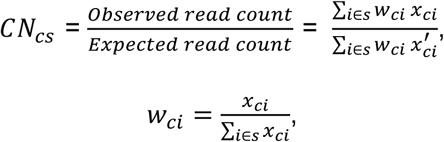

where *c* is a cell, *s* is a segment from DNACopy and it spans across bin *i*, *w*_*ci*_ is the fraction of each bin *i*’s read count out of the total read count of the segment *s*, *x*_*ci*_ is the observed read count, *x*^′^_*ci*_ is the expected normal read count from wavelet transformation and robust PCA normalization.

From DNACopy, if no segmentation is performed, copy number assignment is estimated directly using a negative binomial model. For chromosomes with multiple segments from DNACopy output, copy number is computed using an estimated cell-matched control followed by segmentation.

CHASM is available at: https://github.com/EmiliaCXY/CHASM

### Construction of copy number alteration spike-in datasets

Coassays of scDNA and scATAC from breast epithelial cells preprocessed and copy number analysis on the scDNA modality was performed as described above. Cells with > 99% of the genome with normal copy number state that also passed quality control in scATAC data were retained. A set of 800 cells were chosen randomly as the pool of spike-in cells, and a set of 2000 cells were chosen randomly as the pool of non-spike-in cells. We designed spike-in experiment for all combinations of event size (i.e. full chromosome, chromosome arm, and a 50Mb region), event amplitude (i.e. from two copies loss to two copies gain), and event frequency (i.e. 1%, 5%, and 10%), Each spike-in condition has three replicates. A total of 2000 cells were selected randomly by combining spike-in and non-spike-in cells at a target proportion. For the spike-in cells, read counts from chromosome 1 or chromosome 1p or a fixed 50Mb region in chromosome 1 were removed or added proportionally based on spike-in event amplitude. A cell by 2 Mb genomic bin matrix is constructed for each spike-in condition.

### Copy number analysis of the spike-in data with epiAneufinder

epiAneufinder (v1.0.3) was applied using a window size of 2Mb, with mitochondrial genome excluded in the analysis. This window size is selected to match with the window size used in CHASM. The hg19 genome was used to be consistent with the breast tissue dataset. Blacklist regions were obtained from ENCODE. For the spike-in experiment data, fragment files were generated from the cell by bin read count matrix, by generating fragments of length 60bp with start and end coordinates located within the genomic bins and exporting the resulting per-cell fragment coordinates in fragment file format.

### Copy number analysis of the spike-in data with CopyscAT

CopyscAT (v 0.40) was applied following the tutorial accompanying the original manuscript^20^. Read count matrix of cell by 2Mb genomic bins was generated, and counts were aggregated at the chromosome-arm level within the algorithm. All remaining parameters were set as described in the tutorial. The hg19 genome was used on the breast tissue dataset^17^.

### Evaluation of precision and recall curve from sweeping segmentation thresholds in CHASM and epiAneufinder

In CHASM, the segmentation stringency parameter in DNACopy was varied to assess its effect on downstream copy number assignment. Following initial segmentation with a threshold of 0.005, `segments.p()` was used to calculate p-values of the inferred change points. Segmentation was repeated across 100 p-value thresholds spanning the full range of possible p-values obtained from `segments.p()`.

In epiAneufinder, segment retention was based on the z score of segment-level read depth. To evaluate the effect of this threshold on copy number assignment, segments were refiltered using 100 z-score thresholds spanning the range from −1 to 13, followed by reassignment of copy number states within the algorithm. The range of z-score thresholds were chosen by examining the range of possible z-scores across cells.

Precision and recall were evaluated at a bin level, using 2Mb bins for both CHASM and epiAneufinder. Specifically, 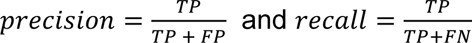. For positive calls, evaluation considered only the direction of copy number change (gain/loss or neutral), and false positives were assessed using the non-spike-in cells. Pareto frontier curves were constructed by sorting results in descending order or recall and retaining the maximum precision at each recall level.

### Evaluation of segmentation localization

Two-sided t-tests were performed on bins within segments surrounding the spike-in breakpoints on chromosome 1p and in the 50 Mb spike-in condition. In the chromosome 1p analysis, bins on chromosome 1p were compared with bins on chromosome 1q. In the 50 Mb spike-in analysis, bins in segments adjacent to the spike-in region were compared with bins within the spike-in segment. For CHASM, statistical testing was based on the signal values obtained after wavelet transformation and robust PCA normalization. For comparison, tests were performed using read-depth values following library-size normalization, square-root transformation, and centering. To examine the effect of wavelet transformation, residual values derived from robust PCA applied directly to the normalized read-depth matrix were also assessed.

### Kidney scMultiome dataset and processing

Nine kidney single-cell multiome samples were obtained from Wilson et al^27^ (GSE232222). For quality control, fragments overlapped with ENCODE blacklist regions were removed, and cells with fewer than 3,000 fragments and TSS enrichment score lower than 4 were removed. Subsequently, fragments were bin into 5 Mb tiles across the hg38 reference genome to produce a cell by bin read count matrix.

### Kidney tissue scRNA transcript read depth summary

For comparison between scATAC inferred copy number alteration and scRNA transcript level, gene counts were aggregated by chromosomal location to generate a cell by chromosome count matrix. The read counts were normalized by library size, square root transformed and scaled across cells.

### p53 knock out mouse pan-tissue dataset and processing

Single-cell ATAC-seq fragment files from the Wang et al. dataset (GSE217661)^32^ were obtained for six mice where three samples were Trp53 wild type and three were Trp53-/-. For quality control, fragments overlapped with ENCODE blacklist regions were removed, and cells with fewer than 3,000 fragments and a median fragment-to-nearest-TSS distance higher than 30 kb were removed. To annotate cell type, gene activity scores were calculated on the scATAC-seq data and used for MaxFuse^72^ to perform label transfer from a mouse scRNA-seq reference (Tabula Muris Senis droplet). Cells flagged as low-confidence by MaxFuse were excluded from downstream analysis. Subsequently, fragments were bin into 2 Mb tiles across the mm10 reference genome to produce a cell by bin read count matrix. Cells were grouped by MaxFuse-assigned lineage, and a lineage was retained in downstream copy number analysis only if both genotypes had at least 100 cells after quality control, yielding six qualifying lineages: thymus antigen presenting cells, pancreas endocrine, liver hepatocyte, and immune cells (myeloid, T, and B cells from across tissues). In this dataset, CHASM was run per lineage, pooling all cells of that lineage across mice, as the immune lineages were under-represented per-sample. epiAneufinder and CopyscAT were applied as outlined in their respective tool tutorials.

### Mouse hematopoietic stem cell dataset and processing

Single-cell ATAC-seq data for young and aged hematopoietic stem cells (HSCs) were obtained from Itokawa et al. (GSE190424)^34^. For quality control, fragments overlapped with ENCODE blacklist regions were removed, and cells with fewer than 3,000 fragments were also removed. Subsequently, fragments were bin into 2 Mb tiles across the mm10 reference genome to produce a cell by bin read count matrix.

### Mouse aging atlas data and processing

Processed single-cell chromatin accessibility data from across 21 mouse tissues at three ages (1, 5, and 21 months) were obtained from Lu et al^26^. Specifically, cell by peak count matrices and for each tissue and associated metadata were downloaded from GSE288730. For quality control, fragments overlapped with ENCODE blacklist regions were removed, and cells with fewer than 3,000 fragments and with fraction of reads overlapping promoter region smaller than 0.25 were also removed. Subsequently, peaks were bin into 5 Mb tiles across the mm10 reference genome to produce a cell by bin read count matrix. The matrices were converted into 10x MTX format for running epiAneufinder and CopyscAT.

To assess age-related trends in CNA burden, the average fraction of genome with CNAs was calculated for each sample and cell type. Within each sample, cell types represented by fewer than 50 cells were excluded. Cell types represented by fewer than two samples per age group and sex were also excluded from the analysis. For each cell type and sex, the association between sample-level CNA burden and age was then tested via Spearman correlation, and Benjamini-Hochberg procedure was applied for multiple testing correction.

### Kidney tissue scATAC atlas preprocessing and annotation

Ninety-nine publicly available human kidney single-cell datasets comprising single-nucleus multiome (RNA + ATAC) and snATAC-seq–only libraries were aggregated to generate an integrated human kidney atlas^27,40–44^. scATAC-seq fragment files were processed with SnapATAC2^73^ using the hg38 reference genome. Cells were filtered based on fragment count, transcription start site enrichment, and doublet status via Scrublet^74^ and DoubletDetection^75^.

For multiome samples, RNA count matrices were filtered based on transcript count and doublet status. ATAC-only datasets were padded with zero-valued RNA features to enable simultaneous processing with multiome data. The combined dataset was integrated using MultiVI^76^, implemented in scvi-tools, and the learned latent representation was used for neighborhood graph construction, UMAP visualization, and clustering. Cell types were annotated based on marker genes. Code used to generate the human kidney atlas is available at https://github.com/GaryWang7/Wang_RAGE24_2026.

### Identifying differentially expressed genes

To identify genes differentially expressed in cells with a specific chromosomal copy number alteration, metacells were constructed from gene expression data by aggregating reads across cells of the same cell type harboring a gain or loss of a given chromosome. For each sample, equal number of cells from the same cell type without any alteration were randomly selected to generate a matched normal metacell profile. Differential expression analysis was performed using DESeq2^77^, with sample identity and chromosomal alteration type included as covariates.

### Copy number profile characterization in kidney cancers

Renal cancer copy number alterations data mapped to the hg19 genome from TCGA were downloaded from the Broad Institute Fire Browse portal (http://gdac.broadinstitute.org). In total 1059 and 593 number of patients were included in clear cell renal cell carcinoma and papillary renal cell carcinoma. Copy gains are defined as log segment ratio greater than 0.3, and copy losses are defined as log segment ratio smaller than −0.3. The copy number alteration frequency was calculated as the fraction of the total patients with gain or loss in the given segment.

### Meta-analysis on genes and transcription factor features associated with mCA burden

For each sample from the kidney single-cell atlas with RNA modality available, SAVER (v1.1.2)^78^ was used to rescue gene expression to address data sparsity. Spearman correlations were then calculated between recovered gene expression and the proportion of the genome affected by copy number alteration across cells of the same type within each sample. Fisher’s z-transformation was applied to the Spearman correlation coefficients to stabilize variance. Transformed coefficients were combined across samples by calculating a weighted average, using the number of cells in each sample as weights, to obtain an overall correlation value. Statistical significance was assessed by z-test, in which the combined z statistic was compared to its standard error.

Genes were subsequently ranked by the combined correlation coefficients, and gene set enrichment analysis was performed using GSEA from the clusterProfiler package (v4.12.6)^79^ with the Hallmark pathways to identify significantly enriched pathways.

